# Gβγ engages PLCβ3 at multiple sites to reorient and facilitate its activation

**DOI:** 10.64898/2026.01.14.699417

**Authors:** Isaac J. Fisher, Kanishka Senarath, Kennedy Outlaw, Kaushik Muralidharan, Elisabeth E. Garland-Kuntz, Michelle Van Camp, Tommy Komay, Leon P. Laskowski, Asuka Inoue, Evi Kostenis, Nevin A. Lambert, Angeline M. Lyon

## Abstract

Phospholipase C β (PLCβ) enzymes are activated by heterotrimeric G protein subunits, increasing hydrolysis of phosphatidylinositol-4,5-bisphosphate (PI(4,5)P2) at the plasma membrane. All four human PLCβ isoforms (PLCβ1-4) are activated by Gα_q_, while PLCβ1-3 are activated to varying extents by Gβγ. The binding sites for Gα_q_ on PLCβ are well-established and much has been learned about its mechanism of activation, but comparatively little is known about Gβγ-dependent activation. In this work, we used cryo-electron microscopy (cryo-EM) single particle analysis (SPA), functional assays, and bioluminescence resonance energy transfer (BRET) to investigate how Gβγ interacts with PLCβ3 in concert with activated Gα_q_ to regulate phospholipase activity. Gβγ heterodimers bind multiple surfaces of PLCβ3 to promote activation but alone do not recruit the enzyme to the plasma membrane. Instead, Gβγ facilitates activation by Gα_q_, most likely by reorienting the phospholipase catalytic site at the membrane to maximize PI(4,5)P2 hydrolysis and downstream Ca^2+^ release. Cell-based functional assays demonstrate that Gβγ is required for maximal PLCβ3 activation even when G_q_ heterotrimers are the sole source of Gβγ. Together, these findings demonstrate that Gβγ acts as a critical positive allosteric modulator that regularly acts in concert with Gα_q_ to activate PLCβ3 at the plasma membrane.

## Introduction

Heterotrimeric G proteins regulate a wide variety of effectors downstream of G protein-coupled receptors (GPCRs). One family of effector enzymes are the phospholipase Cβ (PLCβ) enzymes. The four PLCβ isoforms (PLCβ1-4) cleave phosphatidylinositol-4,5-bisphosphate (PI(4,5)P2) to inositol-1,4,5-trisphosphate (IP3) and diacylglycerol (DAG). These second messengers in turn increase intracellular Ca^2+^ and activate protein kinase C (PKC). All PLCβ isoforms are activated by direct binding of Gα_q_, released by G_q_-coupled receptors. PLCβ2, PLCβ3, and to a lesser extent PLCβ1, are also stimulated by binding of Gβγ heterodimers, released by G_i_-coupled receptors (1, 2).

PLCβs share four core domains with other PLC enzymes, including a pleckstrin homology (PH) domain, four EF hands, a catalytic triose phosphate isomerase (TIM) barrel split by a regulatory linker (X–Y linker) into X and Y subdomains, and a C2 domain. The PLCβ subfamily is defined by its unique proximal and distal C-terminal domains (CTDs) that follow the C2 domain. The proximal CTD (pCTD) includes the autoinhibitory the Hα2*′* helix, which is displaced when Gα_q_ binds (3), and the distal CTD (dCTD), which contributes to membrane binding and contains a second functionally critical Gα_q_ binding site (2). The mechanism by which Gα_q_ binds to and activates PLCβ has been well characterized through functional and structural studies (3–6), but the mechanism by which Gβγ activates PLCβ is much less clear.

The Gβγ heterodimer has no intrinsic enzymatic activity yet regulates a wide variety of effector enzymes via membrane recruitment and/or allostery (7, 8). Prior studies of Gβγ regulation of PLCβ identified two potential, non-overlapping binding sites for Gβγ. The first was the PH domain, which serves primarily as a protein-protein interaction site in the PLCβ subfamily (9, 10). The PH domain is required for Gβγ stimulation of PLCβ and chimeras of PLCδ that contained the PH domain of PLCβ gained sensitivity to the G protein subunit (11). The second Gβγ binding site was mapped to a helix in the Y subdomain of the TIM barrel. Peptides corresponding to this region blocked Gβγ activation of PLCβ and crosslinked to Gβγ (12, 13). However, Gβγ binding to this site appeared to preclude the PLCβ active site from engaging the membrane for PI(4,5)P2 hydrolysis. The mechanism of Gβγ-dependent activation is further complicated by reports that, in cells, the process requires prior or coincident activation by Gα_q_ (14–16).

Recent structural work has shed light onto how Gβγ interacts with PLCβ. Cryo-electron microscopy (cryo-EM) reconstructions of Gβγ and PLCβ on liposomes and nanodiscs showed two Gβγ molecules bound to one PLCβ3, one to the PH domain and one to the EF hand domain (17). The latter domain had not previously been implicated in Gβγ-dependent activation. Gβγ binding did not induce conformational changes within the lipase, and despite its proximity to a lipid bilayer, PLCβ3 remained in its autoinhibited conformation; the active site remained occluded by the X–Y linker and inhibitory interactions between the Hα2*′* helix and the catalytic core persisted. Because no allosteric changes in the lipase were observed, a model was proposed in which Gβγ activates PLCβ3 by recruiting it to and orienting it at the plasma membrane (17). Consistent with this idea, the same study demonstrated Gβγ-dependent partitioning of PLCβ3 to a lipid bilayer (17), although several prior studies using purified components did not show Gβγ-dependent recruitment of the lipase to membranes (18–20). However, the Gβγ–PLCβ3 interface(s) responsible for activation in the cellular environment are unknown. Finally, it is also unclear if G protein activation liberates sufficient free Gβγ to recruit PLCβ3 in cells.

Here we investigate the mechanism by which Gβγ binding to PLCβ3 increases lipase activity using crosslinking, cryo-EM single particle analysis (SPA), and functional assays in living cells. We report cryo-EM reconstructions of Gβγ–PLCβ3 that reveal a third binding site for Gβγ involving the PH, EF hand, and C2 domains. We show that all three structurally determined interfaces contribute to Gβγ-dependent activation in cells. Notably, we find that free Gβγ does not promote recruitment of PLCβ3 to the plasma membrane, and that membrane-tethered PLCβ3 can still be activated by Gβγ. Thus, Gβγ binds to multiple sites on the lipase to further stimulate PI(4,5)P2 hydrolysis concomitant with activation of PLCβ3 by Gα_q_, most likely by orienting the enzyme at the plasma membrane. We propose that Gβγ is best understood as a critical positive allosteric modulator of PLCβ3, as opposed to a bona fide activator in cells.

## Results

### Cryo-EM reconstructions of Gβγ–PLCβ3 complexes in solution

We first attempted to determine the solution structure of a soluble Gβγ–PLCβ3 complex, in which the prenylated Gγ C68 is mutated to serine (Gβγ C68S) (21), eliminating the need for lipids and/or detergents. Complexes of Gβγ C68S–PLCβ3 could be isolated by size exclusion chromatography (SEC) but were too unstable for structural determination, in agreement with previous studies (17). We turned to crosslinking to isolate a stable complex (22). Solvent-exposed cysteines in the lipase (human PLCβ3 residues 193, 221, 358, 516, 824 and 834) were mutated to serines in the background of PLCβ3 Δ892, a C-terminal truncation which lacks the distal CTD yet retains robust activation by Gβγ (23–25). Given the evidence that Gβγ binds to the PH domain, an E60C mutation was installed in the PH domain (PLCβ3 Δ892 PH_cys_) to facilitate crosslinking with Gβγ C68S (Gβ1 contains fourteen cysteines, with C204 and C271 solvent-exposed, and Gγ2 contains two cysteines with only C68 solvent-exposed). As a control, C516 was retained in the X–Y linker (PLCβ3 Δ892 XY_cys_). Neither variant underwent self-crosslinking, in contrast to PLCβ3 Δ892 which retains the endogenous cysteines. Only PLCβ3 Δ892 PH_cys_ crosslinked efficiently to Gβγ C68S (**Figure S1**). Bismaleimidoethane (BMOE), an 8 Å crosslinker had a crosslinking efficiency of >50%, and a 1:1 complex was observed with PLCβ3 Δ892 PH_cys_, consistent with a persistent and specific interaction (**Figure S1B**). To confirm the BMOE-crosslinked Gβγ–PLCβ3 Δ892 PH_cys_ complex was functional, crosslinking was repeated using wild-type Gβγ and PLCβ3 Δ892 PH_cys_, resulting in ∼3-fold greater activity than the reaction without the crosslinker (**Figure S1C**). Similar results were also obtained with the 14.7 Å BM(PEG)2 crosslinker. The crosslinked Gβγ–PLCβ3 Δ892 PH_cys_ complexes were purified using SEC and subjected to cryo-electron microscopy (cryo-EM) single particle analysis (SPA). Two reconstructions were independently refined to 4 Å and 7 Å resolution in the BMOE data set (**Figures 1, S3-S5**, **Table 1**), and one 4.4 Å reconstruction in the BM(PEG)2 data set (**Figures S4-S5, Table 1**).

**Figure 1.**
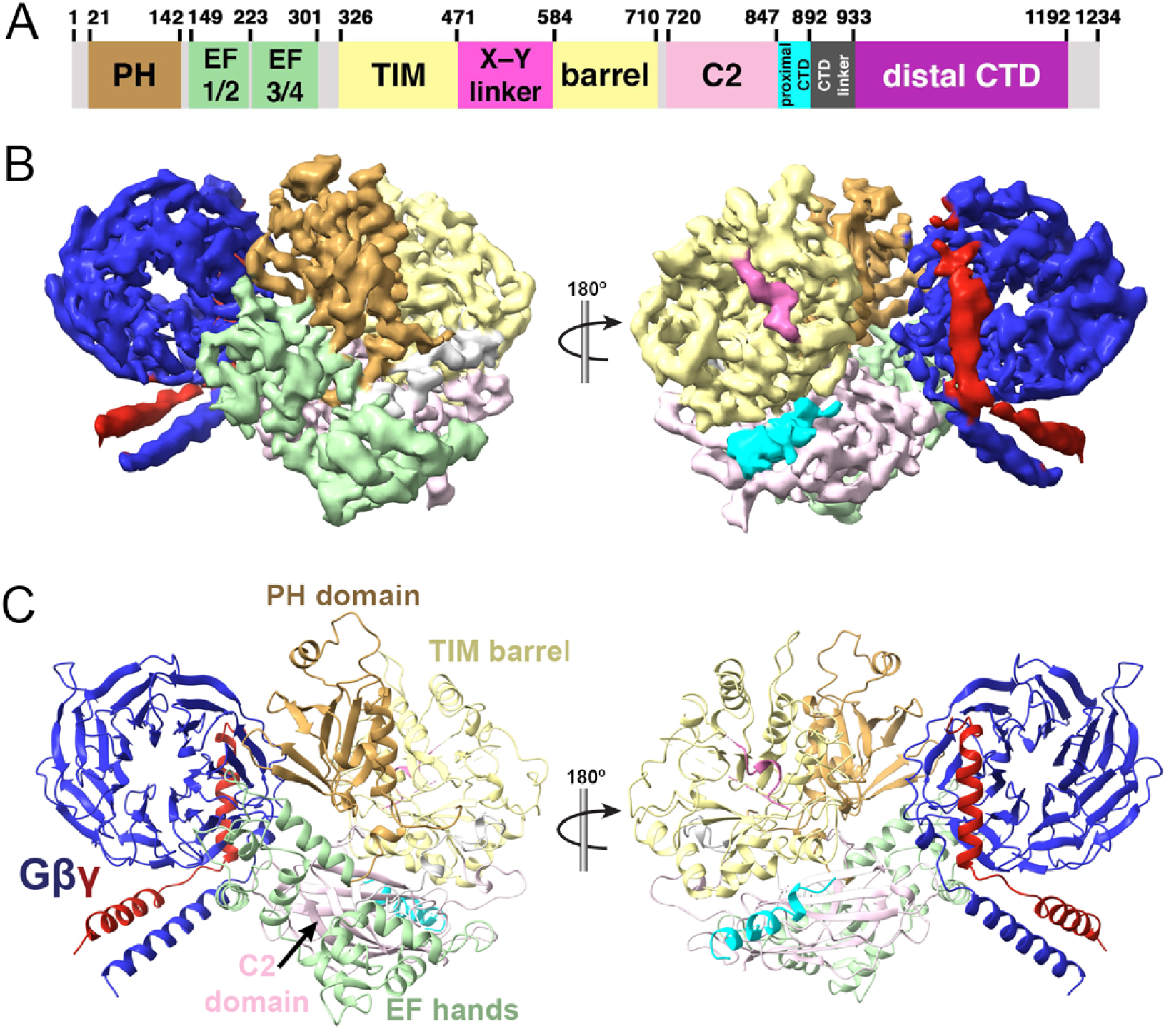
Cryo-EM reconstruction of the Gβγ–PLCβ3 Δ892-PH_cys_ complex. (A) Domain diagram of human PLCβ3, with numbers above corresponding to domain boundaries. PLCβ3 is regulated by the X–Y linker (hot pink), proximal C-terminal domain (pCTD, cyan), and distal CTD (purple). The CTDs are connected by the unconserved CTD linker. (B) Cryo-EM density map and (C) structure of the 4 Å Gβγ–Δ892-PH_cys_ complex crosslinked with BMOE, with PLCβ3 colored as in 1A, Gβ in blue, and Gγ in red.

**Table 1.**
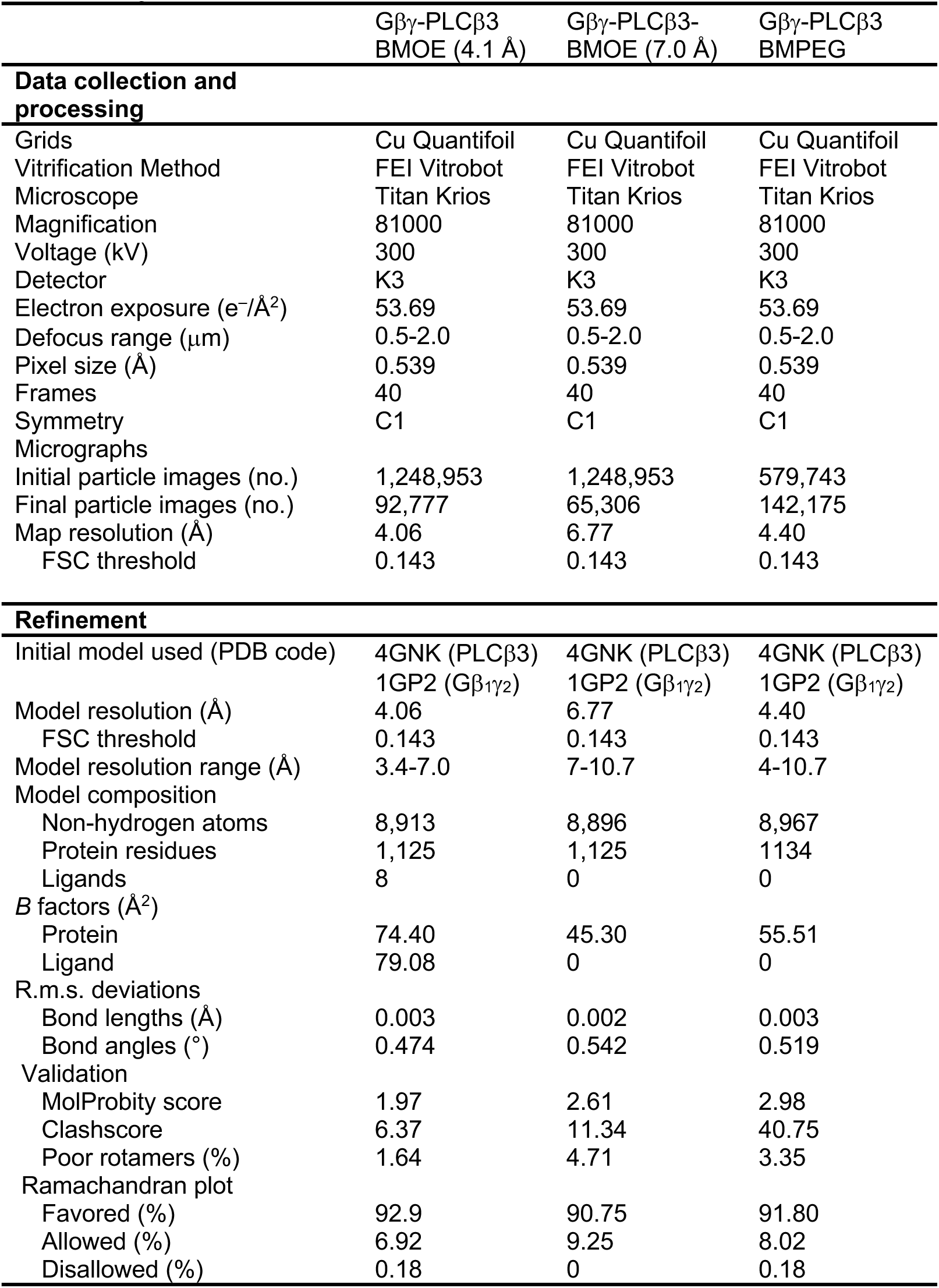
Cryo-EM data collection, refinement and validation statistics.

In all crosslinked Gβγ–PLCβ3 Δ892 PH_cys_ complexes, Gβγ engages PLCβ3 via a surface formed primarily by the PH and EF1/2 domains that differs from the two interfaces observed previously (**Figures S2,3**) (17). The relative orientation and local resolution of Gβγ with respect to PLCβ3 varies in each case, suggesting the interface is conformationally heterogeneous, most likely due to the absence of a membrane (**Figures S2-5**). Even though all the cysteines in Gβ are retained, and both Gβ C271 and C204 are in the “hotspot” interaction surface of Gβγ, only a single crosslink between PLCβ3 E60C and Gβ Cys271 is observed (**Figures S1,3**). In the crosslinked Gβγ–PLCβ3 reconstructions reported, Gβ C204 is ∼25 Å from PLCβ3 E60, well beyond the range of either BMOE or BM(PEG)2. There are also no solvent-exposed cysteines in Gβγ at the EF hand binding site within ∼45 Å of PLCβ3 E60. This is consistent with a specific, persistent interaction that results in the crosslink between Gβ 271 and PLCβ3 E60C. Indeed, density for the crosslinker is observed in the 4 Å BMOE reconstruction, and the orientation of Gβγ and PLCβ3 in the 7 Å BMOE and BM(PEG)2 reconstructions are also consistent with crosslinking via this site.

In the crosslinked complexes, as in the prior structures, PLCβ3 Δ892 PH_cys_ is autoinhibited by its X–Y linker and pCTD (**Figure 1**) (3, 4, 9, 17, 26). This is consistent with previous reports demonstrating the membrane is essential for regulation by Gβγ and that its activation mechanism is independent of the PLCβ CTDs (2). The crosslinked Gβγ is situated such that it allows simultaneous binding of Gα_q_ to PLCβ with the C-terminal helix of Gγ nearly in the same plane as the phospholipase active site. Thus, the solution reconstruction may represent a membrane-localized complex, but not a catalytically active state.

In the 4 Å reconstruction (**Figures 1, 2A**), the Gβγ–PLCβ3 Δ892 PH_cys_ interface buries ∼1,400 Å^2^ surface area. This is more extensive than the Gβγ-PH domain or Gβγ-EF hand interfaces, which bury ∼800 Å^2^ and ∼1,100 Å^2^ respectively (**Figures 2B, 2C**) (17). The primary Gβγ interface is formed by residues on the side of the WD40 toroid, rather than its face, which is the typical effector interface (**Figure S2**). Nevertheless, the crosslinked Gβγ–PLCβ3 Δ892-PH_cys_ interface includes several residues known to be critical for enzyme activation, and the specific interactions differ from the other reported Gβγ–PLCβ3 interfaces (**Figure S6**). In the BMOE complex, Gb D228 interacts with R199 and K183 in PLCb3, whereas in the Gbg–PLCb3–Gbg reconstruction Gb D228 interacts with R24 or K238 in the PH domain and EF hand interfaces, respectively (**Figure 2A-C**). In our Gβγ–PLCβ3 Δ892 PH_cys_ structure, Gβ K301 and R304 on blade 6 make electrostatic interactions with PLCβ3 R741 and E34, respectively (**Figure 2A**). These interactions were not reported in the liposome-bound Gβγ–PLCβ–Gβγ complex (17), and provide a structural explanation for observations reported over thirty years ago by Neer and coworkers on the importance of Gβ blades 6 and 7 in lipase activation (27, 28). On the other hand, Gβ W99, a critical residue for effector activation including PLCβ2 (29), does not interact with the lipase in the crosslinked complexes, but does interact in the Gβγ–PLCβ3–Gβγ complex. Taken together with previous structural findings, our results suggest that Gβγ can bind to PLCβ3 at several sites with modest affinity.

**Figure 2.**
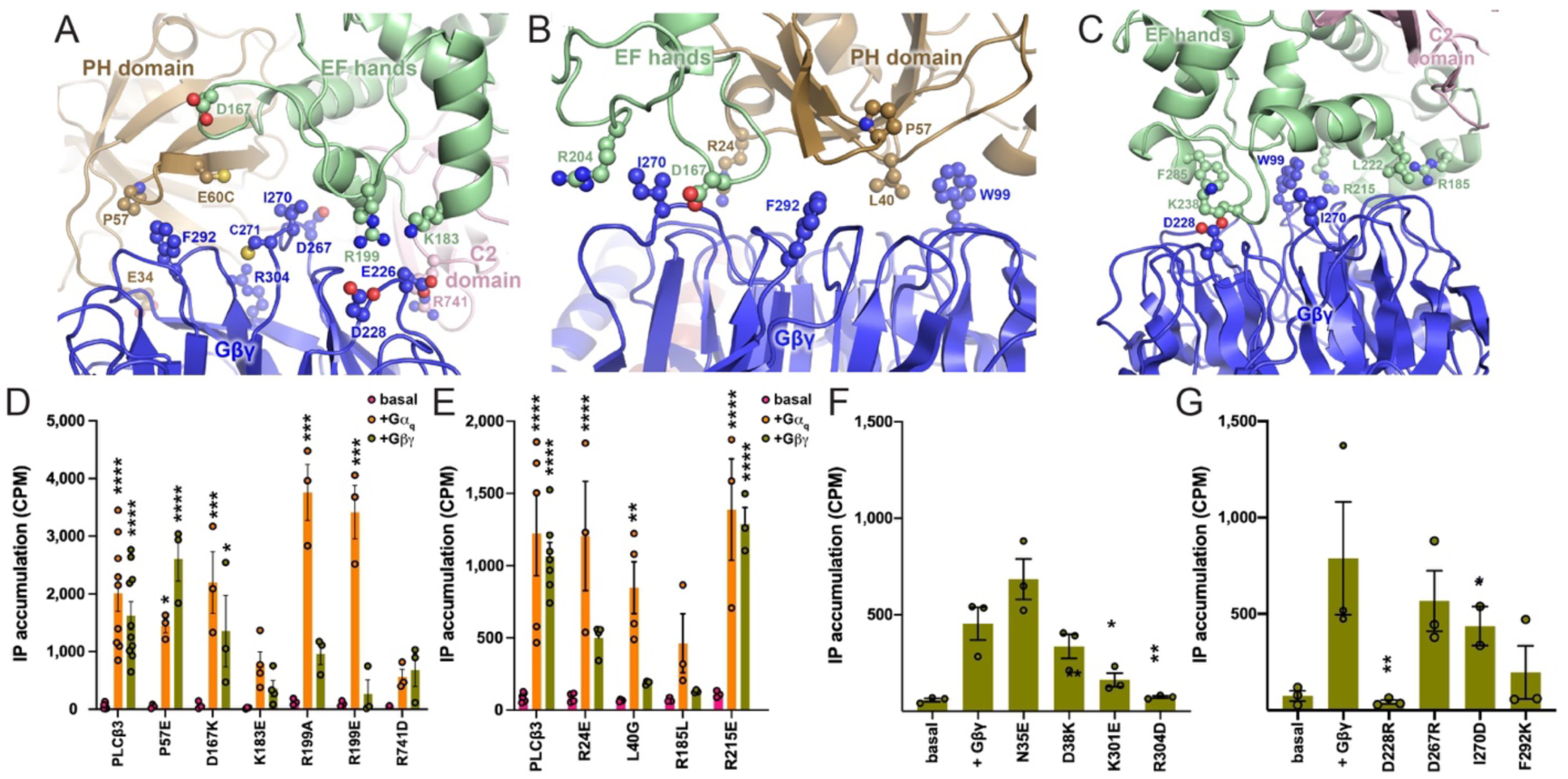
Functional analysis of Gβγ–PLCβ3 interfaces: IP accumulation. Comparison of the (**A**) Gβγ–Δ892-PH_cys_ interface observed in this study, and the previously reported (**B**) Gβγ–PLCβ3 PH and (**C**) Gβγ–PLCβ3 EF hand interfaces (PDB ID: 8EMW) (17). Proteins are colored as in Fig. 1. Residues in PLCβ3 and Gβγ shown as balls and sticks were mutated, and their basal and G protein-dependent activities quantified in a cell-based [^3^H]-IP_x_ accumulation assay. Basal and Gα_q_-dependent activation was used as a control to confirm the PLCβ3 variants were properly folded. Changes in activity are not due to differences in expression (**Figure S7**). The activities of the PLCβ3 mutants in the (**D**) BMOE-crosslinked Gβγ–Δ892-PH_cys_ interface and (**E**) Gβγ–PLCβ3 PH/EF hand interfaces were measured. Mutations to the Gβ_1_ subunit were similarly assessed for (**F**) the crosslinked Gβγ–Δ892-PH_cys_ interface and (**G**) the Gβγ–PH/EF hand interfaces. All assays were performed in triplicate from at least three independent transfections, and data shown are mean ± SEM. Data in D and E were analyzed using a two-way ANOVA followed by Dunnett’s post-hoc multiple comparisons test, comparing the basal activity of each PLCβ3 variant to its activation Gβγ or Gα_q_. ***, p<0.0005, **, p<0.005, *, p<0.05. Data in F and G were analyzed using a one-way ANOVA, followed by Dunnett’s post-hoc multiple comparisons test, comparing each the Gβγ-stimulated activity of each PLCβ3 mutant to that of wild-type PLCβ3. ***, p<0.0005, **, p<0.005, *, p<0.05.

### Functional analysis of Gβγ–PLCβ3 interfaces: IP accumulation

To assess the functional relevance of the Gβγ–PLCβ3 interfaces observed in cryo-EM reconstructions (**Figure 2**), mutations were introduced to maximally disrupt the three observed interfaces by changing amino acid size and/or charge. The activity of the mutants was assessed using a cell-based inositol phosphate (IP) accumulation assays in COS7 cells, where PLCβ3 activity is increased by the overexpression of either Gβγ or Gα_q_. Mutation of residues in PLCβ3 or Gβ_1_ at any one of the three observed interfaces decreased Gβγ-dependent activation of the lipase (**Figures 2D-G**). PLCβ3 mutants with decreased responsiveness to Gβγ are most likely impaired in binding the Gβγ heterodimer, as their expression and activation by Gα_q_ were minimally altered (**Figures 2D-G; Figure S7**). PLCβ3 K183E and R199A, which eliminate electrostatic interactions with Gβ E226 and D228 in the crosslinked complexes, have 3-fold lower Gβγ-stimulated activity, while PLCβ3 R199E was not responsive to Gβγ (**Figures 2D, E**). PLCβ3 R24E, L40G, and R185L, which disrupt interactions with Gβγ observed in the liposome-tethered reconstructions, also showed 3-fold lower activation (**Figure 2E**) (17, 29). Gβ D228R, which interacts with PLCβ3 in all reconstructions (**Figures 2F, G**) (17, 30), decreased activation of PLCβ3 by ∼20-fold, which cannot be fully attributed to reduced protein expression (**Figure S6**). Mutations in Gβ blades 6 and 7, Gβ K301E and R304D, also decreased PLCβ3 activation, in agreement with previous reports (23) (**Figure 2F,G**).

### Functional analysis of Gβγ–PLCβ3 interfaces: PLCβ3 interaction with Gβγ

Under physiological conditions, PLCβ3 is activated in response to GPCR stimulation that activates G_q_ heterotrimers, rather than overexpression of Gα_q_ or Gβγ as in the previous experimental setting. To assess the functional significance of the three Gβγ–PLCβ3 interfaces downstream of receptor activation we developed a BRET assay to monitor Gβγ binding to PLCβ3 in live cells in real time. This effort was complicated by the fact that a large fraction of PLCβ3 is cytosolic, whereas Gβγ is anchored to the plasma membrane. Because of this, recruitment of PLCβ3 to the plasma membrane by any means (e.g. binding to Gα_q_·GTP) would cause an increase in bystander BRET between PLCβ3 and Gβγ that would sum with BRET due to direct interactions between the two. To eliminate this confound, we anchored HiBit-labeled PLCβ3 to the plasma membrane with a C-terminal CAAX motif, which prevents increases in bystander BRET due to translocation and thus isolates signals due to interactions between Venus-Gβγ and the lipase at the plasma membrane. Activation of angiotensin AT_1_ receptors induced a rapid increase in BRET between HiBit-PLCβ3-CAAX (coexpressed with LgBit and Gα_q_) and Venus-Gβγ (**Figure 3A**). This signal was completely blocked by membrane-localized GRK3ct, which binds and sequesters free Gβγ, and enhanced by membrane-localized GRK2RH, which binds and sequesters free Gα_q_·GTP (**Figure 3A**). The latter observation is complementary to our previous finding that sequestering Gβγ enhances Gα_q_ binding to PLCβ3; both findings suggest that PLCβ3 competes with G protein subunits for binding to the complementary G protein subunits (6).

**Figure 3.**
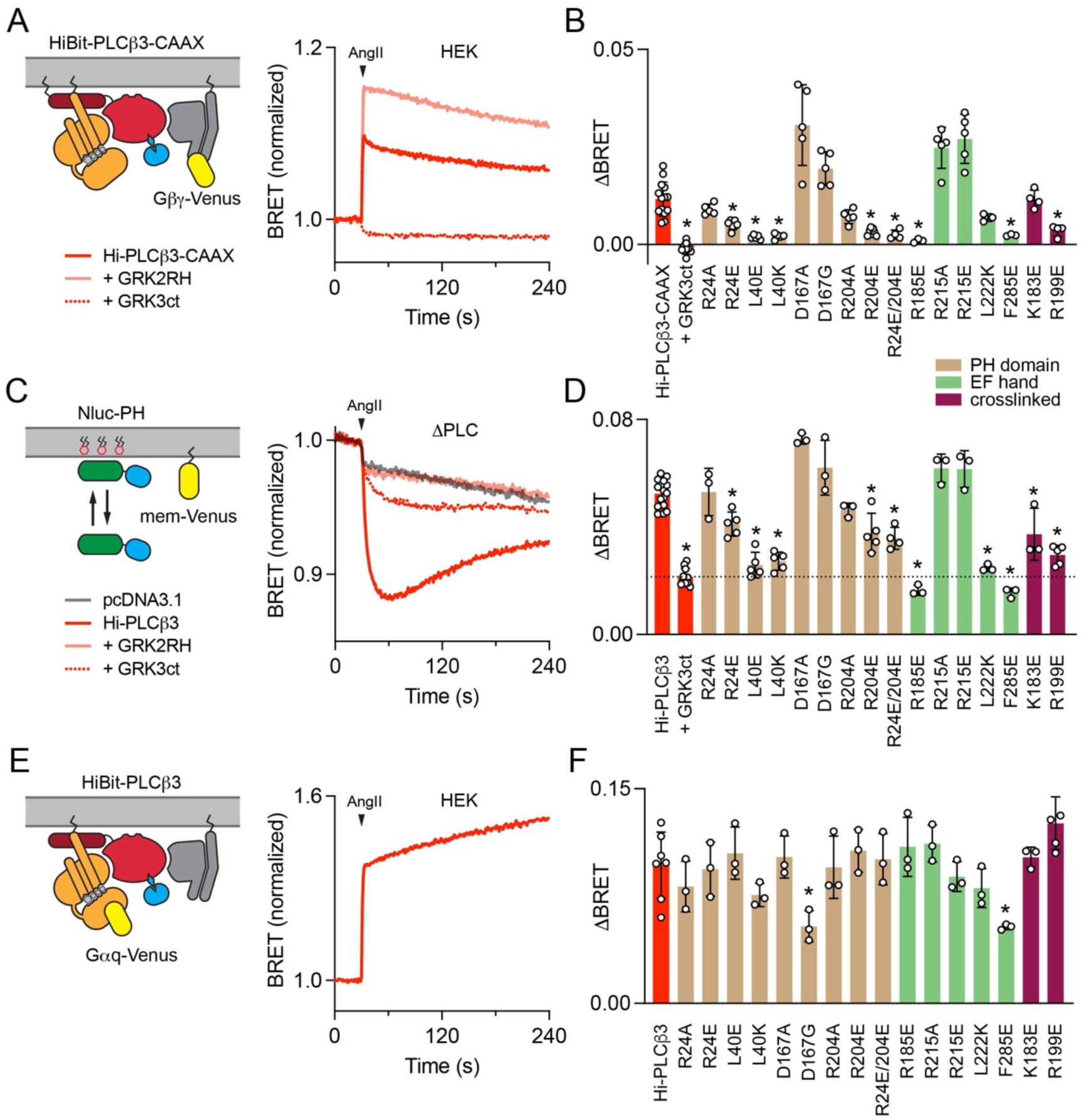
Functional analysis of Gβγ–PLCβ3 interfaces: interaction with G proteins and PI(4,5)P2 hydrolysis. (**A**) BRET between membrane-anchored HiBit-PLCβ3-CAAX and Venus-Gβγ increases after activation of AT_1_ with angiotensin II (AngII; 1 μM). Signals are blocked by membrane-tethered GRK3ct (GRK3ct) and enhanced by membrane-tethered GRK2RH, which sequester free Gβγ and Gα_q_-GTP, respectively. Traces are the average of twenty replicates from five independent experiments. (**B**) AngII-induced changes in BRET between HiBit-PLCβ3-CAAX and Venus-Gβγ (ΔBRET) for sixteen mutants across the three Gβγ–PLCβ3 interfaces; most mutants showed significant changes (denoted by an asterisk) in their interactions with Venus-Gβγ compared to wild-type HiBit-PLCβ3-CAAX. (**C**) In ΔPLC cells, expression of HiBit-PLCβ3 (without LgBit) reconstitutes AngII-induced PI(4,5)P2 hydrolysis, as indicated by bystander BRET between Nluc-PH and mem-link-Venus (mem-Venus). Traces are the average of sixteen replicates from four independent experiments. (**D**) AngII-induced PI(4,5)P2 hydrolysis (ΔBRET) for the same HiBit-PLCβ3 mutants as panel **B**; most mutants showed significant changes in PI(4,5)P2 hydrolysis (denoted by asterisks). (**E**) In HEK cells, BRET between HiBit-PLCβ3 and Gα_q_-Venus increases after activation of AT_1_. Traces are the average of twelve replicates from three independent experiments. (**F**) BRET between the same HiBit-PLCβ3 mutants as panel **B** and Gα_q_-Venus; none of the mutants showed significant changes compared to wild-type HiBit-PLCβ3. For **B**, **D,** and **F**, all mutants were compared to wild-type HiBit-PLCβ3-CAAX or HiBit-PLCβ3 using one-way ANOVA with Dunnett’s post-hoc comparisons; data points represent averages from independent experiments (*n*=3-11) performed in quadruplicate. For all mutants, * p<0.05 and individual p values are given in *SI Appendix*, **Tables S1-5**.

We next used this interaction assay to test a series of sixteen HiBit-PLCβ3-CAAX mutants designed to disrupt the three structurally determined Gβγ–PLCβ3 interfaces by altering amino acid size and/or charge. Mutations in the two interfaces observed in the liposome-tethered reconstructions significantly reduced agonist-induced BRET between Venus-Gβγ and HiBit-PLCβ3-CAAX (**Figure 3B**). For example, L40E in the PH domain interface and R185E in the EF hand interface almost completely abolished receptor-mediated signals (**Figure 3B**). In the crosslinked interface, R199E essentially eliminated the BRET response. The K183E mutant had impaired activation by Gβγ in our IP accumulation assays, but did not significantly reduce agonist-induced BRET signal. Surprisingly, mutations at residues D167 in the PH domain interface and R215 in the EF hand interface significantly *increased* receptor-mediated interaction between Venus-Gβγ and HiBit-PLCβ3-CAAX (**Figure 3B**). Overall, these results support the conclusion that the functional defects observed in our IP accumulation assays reflect loss of Gβγ binding.

Notably, the almost complete loss of agonist-induced BRET signals after disruption of the PH or EF hand interfaces suggests that, when attached to a membrane, Gβγ dimers might cooperate to bind at both sites. This is consistent with the relatively weak binding of Gβγ *in vitro* and difficulties isolating stable Gβγ–PLCβ3 complexes in solution. We then asked if cooperative binding of G protein subunits might extend to Gα_q_, i.e. if binding of Gα_q_ promotes binding of Gβγ. Using the same Gβγ–PLCβ3 interaction assay, we first compared signals downstream of AT_1_, which activates both G_q_ and G_i/o_ heterotrimers, to those downstream of the dopamine D2 receptor (D2R), which activates only G_i/o_ heterotrimers. D2R-mediated signals were detectable, but significantly smaller than AT_1_-mediated signals (**Figure S10A**). Under the same conditions, D2R liberated more free Venus-Gβγ than AT_1_ when detected by the membrane-linked GRK3ct-Nluc sensor (**Figure S10B**), suggesting that the presence of Gα_q_·GTP promotes Gβγ binding to HiBit-PLCβ3-CAAX. To further test this idea we constructed HiBit-PLCβ3-CAAX mutants with disrupted Gα_q_ binding to the proximal CTD (HiBit-PLCβ3-CAAX-LE) or the distal CTD (HiBit-PLCβ3-CAAX-EEE) (6). Interaction of both mutants with Venus-Gβγ was significantly impaired compared to wild-type HiBit-PLCβ3-CAAX (**Figure S10C**). These results suggest that Gβγ binding to the PH and EF interfaces is facilitated by binding of Gα_q_·GTP.

### Functional analysis of Gβγ–PLCβ3 interfaces: receptor-evoked PI(4,5)P2 hydrolysis

To better understand the role of Gβγ binding to different interfaces in receptor-mediated PLCβ3 activation, we carried out PI(4,5)P2 hydrolysis assays based on binding of the PH domain of PLCδ to PI(4,5)P2 at the plasma membrane. PI(4,5)P2 hydrolysis releases Nluc-PH-PLCδ (Nluc-PH) into the cytosol, which is detected as a loss of bystander BRET to a marker on the plasma membrane (mem-Venus; **Figure 3C**). We first established that endogenous Gβγ contributes to activation of endogenous PLCβ3 (the only PLCβ isoform expressed at significant levels in HEK 293 cells) downstream of AT_1_ receptors. Accordingly, PI(4,5)P2 hydrolysis after activation of AT_1_ was inhibited by GRK3ct (**Figure S11A, B**). Because AT_1_ receptors activate both G_q_ and G_i/o_ heterotrimers, we were interested to know the source of the Gβγ contributing to PLCβ3 activation. We found that pertussis toxin (PTX) partially blocked AT_1_-mediated PI(4,5)P2 hydrolysis, consistent with Gβγ from G_i/o_ heterotrimers stimulating lipase activity (**Figure S11A, B**). However, GRK3ct still inhibited PI(4,5)P2 hydrolysis when PTX was present, indicating Gβγ from G_q_ heterotrimers contributes as well (**Figure S11A, B**).

Next, we reconstituted the same PI(4,5)P2 hydrolysis assay in genome-edited HEK 293 cells lacking endogenous PLCβ proteins (ΔPLC cells) by expressing wild-type (wt) or mutant HiBit-PLCβ3. PI(4,5)P2 hydrolysis mediated by expressed HiBit-PLCβ3 was inhibited by GRK3ct to roughly the same extent as that mediated by endogenous PLCβ3 in parental cells (**Figure 3C**). Notably, PI(4,5)P2 hydrolysis under these conditions was completely blocked by GRK2RH, consistent with the idea that activation of PLCβ3 by Gβγ in cells requires Gα_q_·GTP (**Figure 3C**). PI(4,5)P2 hydrolysis signals mediated by HiBit-PLCβ3 variants with impaired Gβγ binding were significantly smaller than signals mediated by wt HiBit-PLCβ3 (**Figure 3D**). Among the most defective variants were the L40E PH domain and R185E EF hand mutants, the latter supporting very modest PI(4,5)P2 hydrolysis comparable to that remaining when Gβγ is sequestered by GRK3ct (**Figure 3D**). Consistent with the Gbg sequestration, these mutants were still significantly impaired in the presence of PTX (**Figure S11C, D**). R199A and K183E, the crosslinked complex interface mutants, also had significantly reduced agonist-induced PI(4,5)P2 hydrolysis (**Figure 3D**). Conversely, in agreement with our Gβγ interaction results, the D167A mutant in the PH domain interface significantly *increased* receptor-mediated PI(4,5)P2 hydrolysis (**Figure 3D**). Overall, there was an excellent correlation between Gbg binding and PI(4,5)P2 hydrolysis across the sixteen mutants we tested, and good agreement with our IP accumulation assays. There were no significant differences between wt HiBit-PLCβ3 and any mutant with respect to association with Gα_q_-Venus (**Figure 3E, F**) or expression level (*SI Appendix,* **Table S1**), ruling out indirect effects on Gα_q_ binding or protein stability. These findings indicate that all three of the structurally resolved Gβγ–PLCβ3 interfaces contribute to activation of PLCβ3 downstream of AT_1_ receptors.

### Gβγ facilitates activation of membrane-anchored PLCβ3

PLCβ3 is not tightly anchored to the plasma membrane, and Gβγ has been proposed to increase lipase activity by recruiting the holoenzyme to the membrane. We previously found that AT_1_ activation results in recruitment of PLCβ3 to the plasma membrane, but concluded that this was primarily mediated by binding to Gα_q_·GTP (6). To test the possibility that Gβγ binding contributes to membrane recruitment of PLCβ3, we used our previously established bystander BRET recruitment assay (**Figure 4A**). We found that none of the mutations that inhibited Venus-Gβγ binding and PI(4,5)P2 hydrolysis significantly impaired recruitment of HiBit-PLCβ3 to the plasma membrane (**Figure 4B**), indicating that recruitment depends entirely on binding to Gα_q_·GTP (6). As an alternative test of this hypothesis, we asked if Gβγ increased the activity of HiBit-PLCβ3-CAAX, which is tightly anchored to the plasma membrane and therefore not subject to translocation from the cytosol. We found that GRK3ct was effective at inhibiting PI(4,5)P2 hydrolysis by HiBit-PLCβ3-CAAX (**Figure 4C**), similar to its effect on PI(4,5)P2 hydrolysis mediated by endogenous PLCβ or HiBit-PLCβ3. Likewise, mutations in HiBit-PLCβ3-CAAX that impaired Gβγ binding also impaired PI(4,5)P2 hydrolysis (**Figure 4C, D**). Taken together, these results suggest that Gβγ does not facilitate PLCβ3 activation by recruiting the lipase to the plasma membrane, but rather enhances the catalytic activity of Gα_q_-bound PLCβ3 through a distinct allosteric mechanism at the membrane.

**Figure 4.**
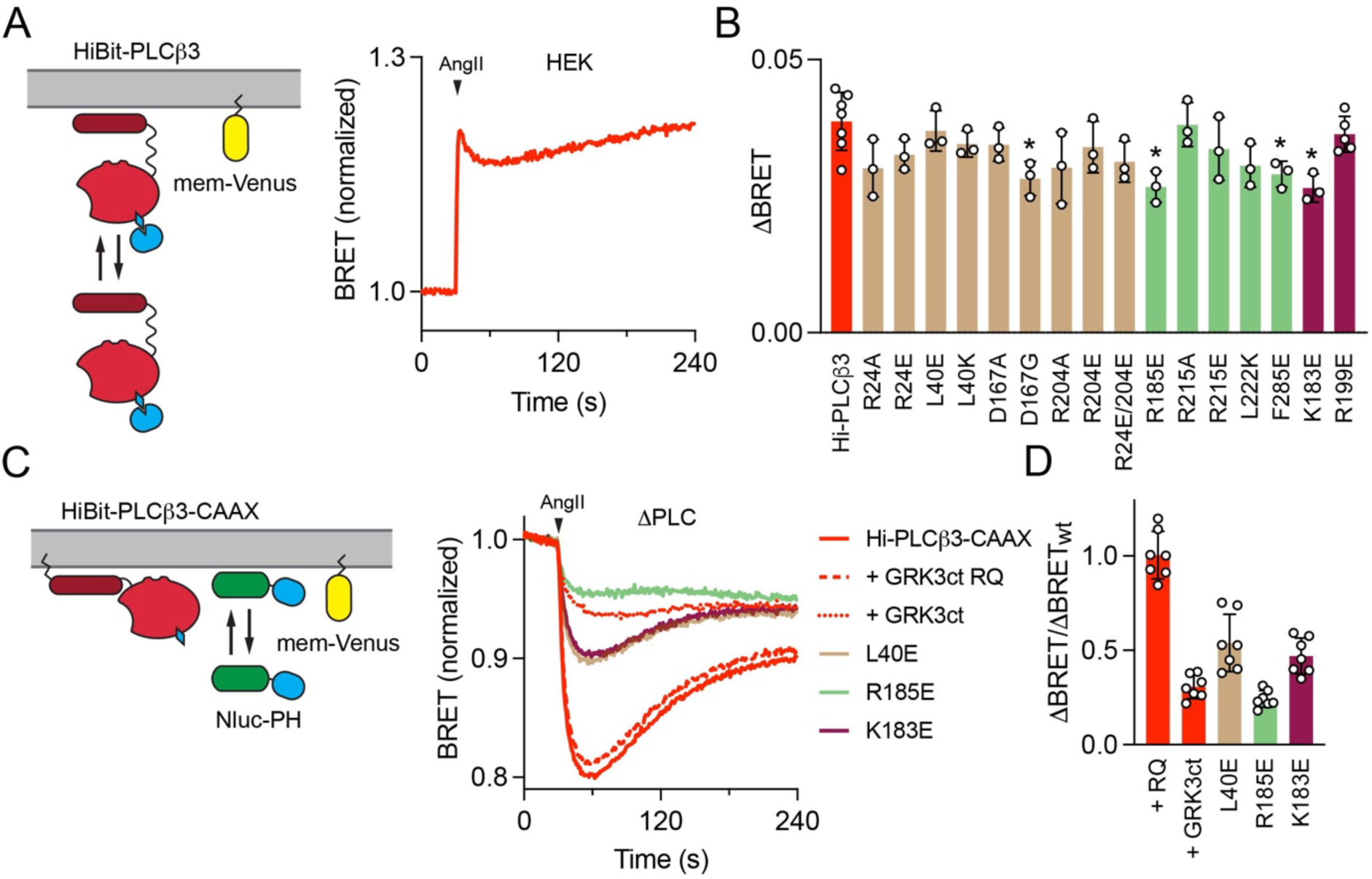
Gβγ facilitation of PLCβ3 activation does not reflect membrane recruitment. (**A**) Activation of AT_1_ induces translocation of HiBit-PLCβ3 to the plasma membrane as indicated by bystander BRET between HiBit-PLCβ3 and mem-Venus. Traces are the average of twelve replicates from three independent experiments. (**B**) BRET between the same HiBit-PLCβ3 mutants as Figure 3 and mem-Venus; none of the mutants showed a significant defect in membrane translocation compared to wild-type HiBit-PLCβ3. Individual p values are given in *SI Appendix*, **Table S6**. (**C**) In ΔPLC cells, expression of HiBit-PLCβ3-CAAX reconstitutes AngII-induced PI(4,5)P2 hydrolysis, indicated by bystander BRET between Nluc-PH and mem-Venus. Signals are inhibited by membrane-tetheredGRK3ct, which sequesters free Gβγ. Traces are the average of twenty-eight replicates from seven independent experiments. (**D**) PLCβ3-CAAX variants shown to be defective with respect to Gβγ binding are also defective with respect to PI(4,5)P2 hydrolysis. For **B,** all mutants were compared to wild-type HiBit-PLCβ3, and for **D** all mutants were compared to +GRK3ct RQ (a mutant defective in binding Gα_q_·GTP), using one-way ANOVA with Dunnett’s post-hoc comparisons; data points represent averages from independent experiments (*n*=3 or 7) performed in quadruplicate. For all mutants, * p<0.05 and individual p values are given in *SI Appendix*, **Tables S6-7**.

## Discussion

PLCβ enzymes have critical roles in numerous processes, from cardiovascular and vascular smooth muscle function to opioid sensitivity (31–35). PLCβ basal activity is tightly controlled, and direct binding of Gα_q_ and Gβγ subunits, released downstream of G_q_- and G_i/o_-coupled receptors, is essential for robust PI(4,5)P2 hydrolysis (2). The mechanisms by which Gα_q_ interacts with PLCβ to increase lipase activity have been well-characterized through structural and functional studies (2, 3, 5, 6, 36). Much less is known about how Gβγ binds to and activates PLCβ1-3. Prior studies established roles for the PH domain and, to a lesser extent, the TIM barrel in Gβγ-dependent activation (11–13, 18, 22, 37). The cryo-EM structure of a liposome-tethered Gβγ–PLCβ3 complex confirmed that Gβγ bound directly to the PH domain, and surprisingly, that a second Gβγ molecule bound to the EF hands (17). However, the functional relevance of both binding sites remained unclear. Moreover, binding of Gβγ also failed to induce conformational changes in the lipase necessary for activation, despite the proximity to a membrane surface. A model was proposed in which Gβγ activates PLCβ3 via membrane recruitment and orientation (17). However, this model contradicted previous studies showing Gβγ did not recruit PLCβ to the plasma membrane (12, 18, 22, 25, 38–40). Interpreting membrane recruitment studies is further complicated by the use of proteoliposome sedimentation assays, the results of which can be confounded by aggregation of proteins on the liposome surface (41), heterogeneity in size, composition, and curvature (42), propensity to phase separate (43), and difficulties in quantifying free versus liposome-bound protein (44). Thus, the question of mechanism remained unsettled.

In this study, we used cryo-EM, activity and signaling assays, and BRET to validate Gβγ–PLCβ3 interactions, assess their contribution to PI(4,5)P2 hydrolysis, and establish whether Gβγ activates PLCβ3 via membrane recruitment. Using cryo-EM, we identified a third Gβγ binding site on the PLCβ3 PH domain and confirmed all structurally determined Gβγ binding sites are necessary for maximum G protein-stimulated activity (**Figures 1-3**). Multiple Gβγ subunits binding to a single effector is not without precedent, as two Gβγ molecules must bind to non-overlapping sites on phosphatidylinositol 3-kinase γ (PI3Kγ) for maximum membrane recruitment and activation (**Figure S6**)(45, 46). The structures of Gβγ–PLCβ3 complexes all tether the lipase to the membrane (**Figure 5**), but it is unlikely that all three sites could be occupied simultaneously during catalysis. In the crosslinked Gβγ–PLCβ3 complex, the orientation of Gβγ is unlikely to be compatible with the binding of the lipase active site to the membrane (**Figures 1, 5A**). We propose this reflects an encounter complex, or pre-activation state, given the specificity and efficiency of the crosslinking reaction (**Figure S1**). In contrast, both Gγ subunits in the liposome-tethered Gβγ–PLCβ3–Gβγ complex and the lipase active site can simultaneously interact with the membrane (**Figure 5B**). Our observation that disruption of either of these sites can severely compromise Gβγ binding suggests a degree of cooperativity between these sites.

**Figure 5.**
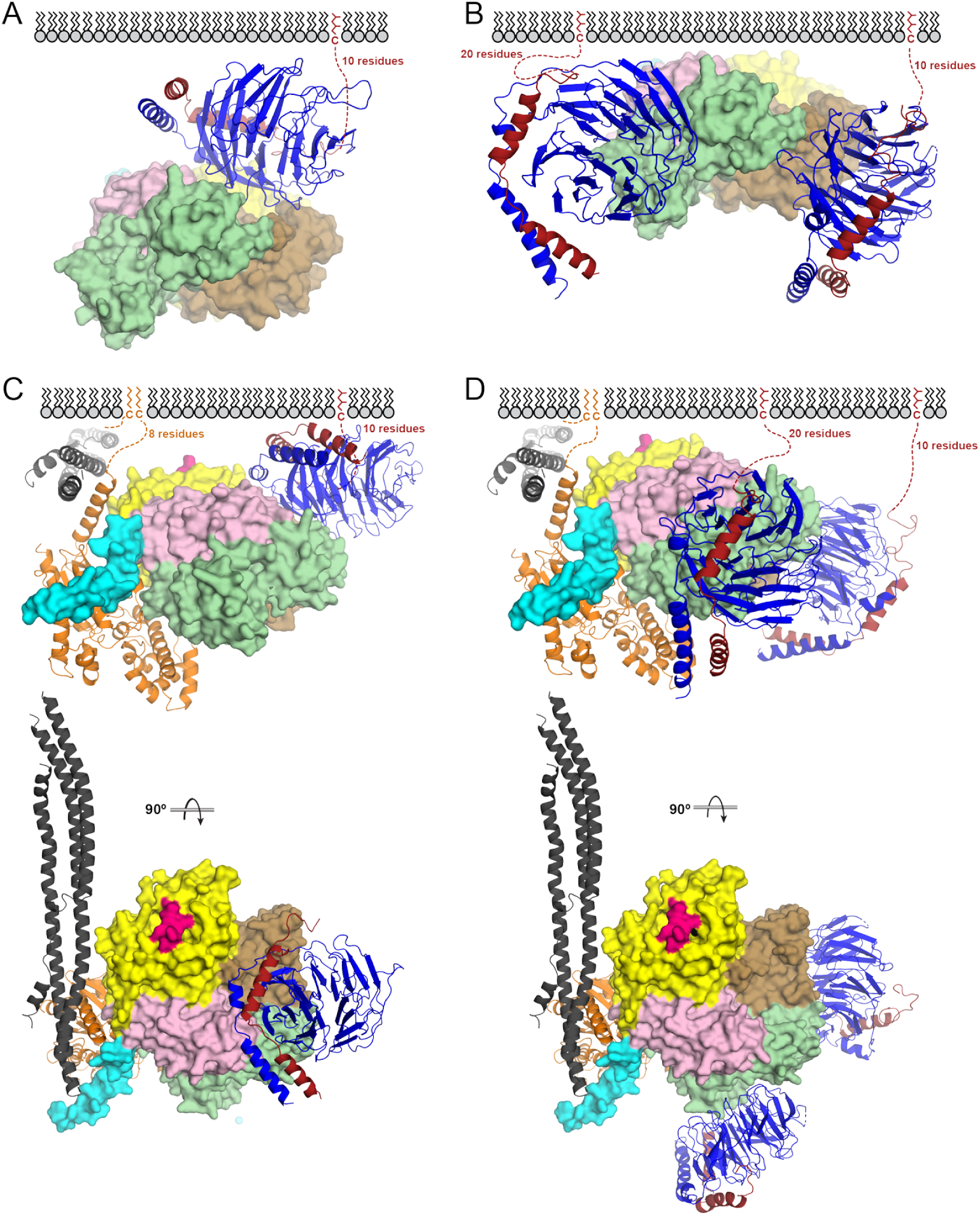
G protein–PLCβ3 complexes at the membrane. (A) The crosslinked Gβγ–PLCβ3 complex is compatible with membrane localization, but not lipase activity. PLCβ3 is shown as a surface, and Gβγ in ribbon. Proteins are colored as in Figure 1A. (**B**) The liposome-tethered Gβγ–PLCβ3 complex would allow the PLCβ3 active site to interact with the membrane. (**C**) The crosslinked and (**D**) liposome-tethered complexes are compatible with Gα_q_·GTP binding and activation via displacement of the Hα2’ helix (cyan) and engagement of the dCTD (dark gray). In both models, the dCTD binds the membrane through electrostatic interactions.

Comparisons of the predicted Gβγ binding sites in PLCβ1-4 provide some insights into the isoform-specific differences in Gβγ-dependent activation. Of the ten residues we identified as functionally relevant across the three PLCb3 Gbg binding sites (**Figures 2, 3**), only two are conserved in PLCb4. The other residues are incompatible with Gβγ binding and likely explain why Gβγ does not activate this isoform (47, 48). In PLCβ1 and PLCβ2, seven of the residues identified are conserved, the exceptions being PLCβ3 R204 and R215. In PLCβ1, these residues are replaced by proline and valine, respectively, and in PLCβ2 they are asparagine and serine. Whether these differences are sufficient to explain why PLCβ1 is weakly activated by Gβγ while PLCβ2 is robustly activated remains to be established (1).

Previous work has shown that PLCβ3 must be preactivated, or simultaneously activated, by Gα_q_·GTP in cells before Gβγ binds to further increase PI(4,5)P2 hydrolysis (14). This is consistent with the fact that all Gβγ–PLCβ3 reconstructions are fully compatible with simultaneous binding Gα_q_·GTP at the membrane (**Figure 5C, D**). Gα_q_ binding is essential for translocation of the lipase to the plasma membrane (6), and mutation of any of the three structurally characterized Gβγ binding sites has no impact on lipase translocation (**Figure 3**). Moreover, free Gβγ released by activation of G_i/o_ heterotrimers does not promote translocation (6). While Gβγ clearly does not increase lipase activity via membrane recruitment, its binding to the lipase is essential for maximum PI(4,5)P2 hydrolysis (**Figure 3**), even when PLCβ3 is tethered at the membrane (**Figure 4**). While Gβγ recruitment and reorientation of PLCβ3 at the membrane were proposed as activation mechanisms, they were not directly tested. Here, we have elucidated this mechanism, and our results demonstrate that Gβγ is a positive allosteric modulator of PLCβ3, following direct activation of the phospholipase by Gα_q_, as opposed to a *bona fide* activator. In this paradigm, Gβγ binding to the PH domain and/or EF hands optimizes the orientation of Gα_q_–PLCβ3 at the membrane to facilitate interfacial activation and maximize PI(4,5)P2 hydrolysis (**Figure 5E**).

Finally, our results show that Gβγ is surprisingly important for PLCβ3 activation by G_q_ heterotrimers in cells. While synergistic activation of PLCβ3 by Gα_q_ and Gβγ is well known, this synergy is typically discussed as a mechanism of crosstalk between G_q_ and G_i_ signaling, the latter being the source of Gβγ dimers (49, 50). Gβγ affinity for PLCβ3 is relatively low and only G_i/o_ heterotrimers are thought to be expressed at high enough levels to release sufficient free Gβγ. Here we show that PI(4,5)P2 hydrolysis downstream of G_q_ alone is sensitive to sequestration of Gβγ dimers, consistent with a previous study in HeLa cells (51), as well as mutations that disrupt Gβγ binding. Endogenous G_q_ heterotrimers are evidently capable of releasing sufficient free Gβγ to occupy binding sites on PLCβ3. This is most likely enabled by cooperative binding of G protein subunits to PLCβ3 at the plasma membrane.

## Materials and Methods

### Cell culture and transfection

Human embryonic kidney HEK 293 cells were obtained from ATTC (CRL-1573) and used as supplied or after gene editing as described previously (15, 52); ΔPLCβ1-4 (ΔPLC) cells were generated using CRISPR/Cas9 and validated as described previously (15). Cells were transfected in 6-well plates in growth medium using linear polyethyleneimine MAX (Polysciences) at a nitrogen/phosphate ratio of 20 and were used for experiments 24-48 hours later. Up to 3.0 μg of plasmid DNA was transfected in each well of a 6-well plate. COS-7 cells were a gift from A.V. Smrcka and used for [^3^H]-IP_x_ accumulation assays.

### Plasmids

SNAPf-AGTR1, HiBit-PLCβ3, HiBit-PLCβ3-CAAX, mem-link-Venus, mem-GRK3ct, mem-link-amber-GRK2RH, Nluc-PH, Gαq-Venus, CMV-LgBit, Venus-1-155-Gγ2 and Venus-156-239-Gβ1 were described previously (6). Mutations in HiBit-PLCβ3, HiBit-PLCβ3-CAAX were made by amplifying three fragments from wild-type plasmids with the desired mutation incorporated in a primer, assembling using Gibson assembly and verifying by full plasmid sequencing. Mutations in human PLCβ3 in pFastbac1, pCMV, or pCDNA3.1 were generated using the Takara infusion site-directed mutagenesis kit (Takara) and verified by full plasmid sequencing. The same strategy was used to subclone Gβ_1_ and Gγ_2_ in pCI-Neo(53) and Gα_q_ in pCDNA3.1+.

### Protein expression, purification, and complex formation

#### Protein expression

PLCβ3, PLCβ3 Δ892 and variants, Gβγ, and Gβγ C68S were expressed and purified from baculovirus-infected insect cells. Baculoviruses were generated using the FastBac recombinant baculovirus system (Invitrogen/Thermo Fisher Scientific, Inc.) in ESF 921 Insect Cell Culture Medium (Expression Systems)-adapted Sf9 (*Spodoptera frugiperda*) cells.

#### Purification of Gβγ and Gβγ C68S

High Five cells were infected with baculoviruses encoding His6-Gα_i1_, Gβ_1_, and Gγ_2_ (54, 55). Cells were harvested 60 h post-infection by centrifugation at 2,500 x *g*, frozen in liquid N_2_ and stored at −80 °C.

All purification steps were performed at 4 °C unless otherwise indicated. Cell pellets were thawed in 15 mL of lysis buffer (50 mM HEPES pH 8.0, 3 mM MgCl_2_, 10 mM β-mercaptoethanol, 0.1 mM EDTA, 100 mM NaCl, 10 μM GDP, and protease inhibitors (133 μM PMSF, 21 μg/mL TLCK, and 0.5 μg/mL TPCK) and lysed by four freeze-thaw cycles in liquid nitrogen. Lysate was diluted to 100 mL with lysis buffer and centrifuged at 100,000 x *g* for 30 min to isolate the membrane fraction. The membrane pellets were resuspended by dounce in 5 mL extraction buffer (50 mM HEPES pH 8.0, 3 mM MgCl_2_, 50 mM NaCl, 10 mM β-mercaptoethanol, 10 μM GDP, and protease inhibitors), combined and diluted to 60 mL. Cholate was added to a final concentration of 1% and the mixture stirred for 1 h at 4 °C to extract membrane proteins. Detergent extracts were clarified by centrifugation at 100,000 x *g* for 45 min. The supernatant was diluted five-fold with buffer A (50 mM HEPES pH 8.0, 3 mM MgCl_2_, 10 mM β-mercaptoethanol, 100 mM NaCl, 10 μM GDP, 0.5% polyoxyethylene(10) lauryl ether (C12E10), and protease inhibitors) and applied to a cOmplete™ His-Tag Purification Resin (Roche) column pre-equilibrated with buffer A. The column was washed with 100 mL of buffer A supplemented with 300 mM NaCl and 5 mM imidazole, then transferred to room temperature and washed with 12 mL buffer A. Gβ_1_γ_2_ subunits were released from His6-Gα_i1_ using six 4 mL fractions of RT buffer (buffer A supplemented with 150 mM NaCl, 5 mM imidazole, 50 mM MgCl_2_, 10 mM NaF, 10 μM AlCl_3_, and 1% cholate). Fractions were analyzed by SDS-PAGE and Coomassie staining to assess purity. Fractions containing Gβγ were pooled and applied to a MonoQ column pre-equilibrated with 20 mM HEPES, pH 8, 1 mM DTT, 50 mM NaCl and 0.5% CHAPS, and eluted with a 50-500 mM NaCl gradient. Fractions containing purified protein were identified by SDS-PAGE, concentrated to 20-40 μM, flash frozen in liquid N_2_, and stored at −80 °C.

Gβγ C68S was purified as described with some modifications. Briefly, after cell lysis, the supernatant was diluted five-fold with buffer A lacking polyoxyethylene(10) lauryl ether, glass fiber-filtered, and applied to cOmplete™ His-Tag Purification Resin (Roche) as described. The rest of the steps were carried out as described, with the omission of cholate or CHAPS from the buffers.

#### Purification of PLCβ3 and variants

His6-PLCβ3, PLCβ3 Δ892, PLCβ3 Δ892-PH_cys_ and PLCβ3 Δ892-XY_cys_ were expressed in Sf9 cells grown in S6900 II serum-free media and infected with baculovirus at an MOI of 1 (56). After 48 h, cells were harvested by centrifugation, frozen in liquid N_2_ and stored at −80°C. Cells were rapidly thawed and lysed by four freeze-thaw cycles in liquid nitrogen in lysis buffer (20 mM HEPES, pH 8, 50 mM NaCl, 10 mM β-mercaptoethanol, 0.1 mM EDTA, 0.1 M EGTA, 133 μM PMSF, 21 μg/ml TLCK and TPCK, 0.5 μg/ml aprotinin, 0.2 μg/ml Leupeptin, 1 μg/ml Pepstatin A, 42 μg/ml Tosyl-L-Arginine Methyl Ester (TAME), 10 μg/ml Soy Bean Trypsin Inhibitor (SBTI)) Lysed cells were collected and diluted with lysis buffer and NaCl to a final concentration of 800 mM NaCl, and centrifuged at 100,000 × *g* for 1 h. The supernatant was diluted five-fold with lysis buffer containing 0.5% polyoxyethylene(10) lauryl ether (C12E10) and centrifuged again at 100,000 × *g* for 1 h The supernatant was loaded onto a cOmplete™ His-Tag Purification Resin (Roche) column pre-equilibrated with buffer A (20 mM HEPES, pH 8, 100 mM NaCl, 10 mM β-mercaptoethanol, 0.1 mM EDTA, and 0.1 M EGTA). The column was washed with 3 column volumes (CVs) of buffer A, followed by 3 CVs of buffer A supplemented with 300 mM NaCl and 10 mM imidazole. The protein was eluted with 3-10 CVs of buffer A supplemented with 200 mM imidazole. Proteins were concentrated and loaded onto tandem Superdex 200 columns (10/300 GL; GE Healthcare) equilibrated with SEC buffer (20 mM HEPES pH 8, 200 mM NaCl, 2 mM DTT, 0.1 mM EDTA, and 0.1 M EGTA). Fractions containing purified protein were identified by SDS-PAGE and were pooled, concentrated, and flash frozen in liquid N_2_.

#### Crosslinking and complex isolatio

Purified Gβγ-C68S, Gβγ, PLCβ3 Δ892, PLCβ3 Δ892-PH_cys_, and PLCβ3 Δ892-XY_cys_ were buffer exchanged to remove DTT by concentrating the proteins in an Amicon Ultra 0.5 ml 30 K concentrator (Millipore-Sigma) and washing them twice with 20 mM HEPES pH 7.4, 100 mM NaCl, 0.1 mM EDTA and 0.1 mM EGTA (and 0.5% CHAPS for Gβγ). 25 μM of the buffer-exchanged Gβγ and 25 μM PLCβ3 Δ892, PLCβ3 Δ892-PH_cys_ or PLCβ3 Δ892-XY_cys_ were mixed and crosslinking initiated by addition of 200 μM BMOE or BM(PEG)2. Reactions were incubated for 45 min at room temperature and quenched by addition of 20 mM DTT. Crosslinking of Gβγ and PLCβ3 Δ892-PH_cys_ were completed as above but contained 0.5% CHAPS. Crosslinking was confirmed by SDS-PAGE by the presence of a band at ∼135 kDa, consistent with a 1:1 stoichiometric complex (Gβ MW: 35 kDa, PLCβ3-Δ892 MW: 100.89 kDa).

### Cryo-EM

#### Sample preparation and data collection

For the BMOE-crosslinked and BM(PEG)2-crosslinked Gβγ C68S–PLCβ3 Δ892-PH_cys_ complexes, 3.5 μL of purified complex at 1 mg/mL supplemented with 0.2 % CHAPS_(f)_ ∼5 min before blotting was applied onto glow-discharged Quantifoil R1.2/1.3 300-mesh grids and prepared and imaged as described for PLCβ3. For the BMOE sample, micrographs were collected on a Titan Krios G1 electron microscope (FEI) equipped with a post-GIF K3 direct electron detector (Gatan) in the Purdue Life Sciences Cryo-EM Facility. A dataset containing ∼4,515 images was collected in super-resolution mode with a pixel size of 0.539 Å, at a defocus range of 1-3 μm using Leginon. For each movie stack, 40 frames were recorded at a frame rate of 78 ms per frame and a total dose of 53.69 electrons/Å^2^. For the BM(PEG)2 sample, micrographs were collected on a Titan Krios G4 electron microscope (FEI) equipped with a Post-GIF K3 direct electron detector (Gatan) in the Purdue Life Science Cryo-EM Facility. A dataset containing ∼4,587 images was collected in super-resolution mode with a pixel size of 0.539 Å, at a defocus range of 1-3 μm using EPU. For each movie stack, 40 frames were recorded at a frame rate of 78 ms per frame and a total dose of 53.43 electrons/Å^2^.

#### Data processing

Micrographs were motion aligned and motion corrected using motioncor2 (57) implemented within CryoSPARC (58). CTF estimations were completed using CTFfind4 (59). Particle picking, 2D classifications, initial model generation, 3D classification and refinement were all performed using CryoSPARC. Workflows for each data set are shown in **Figure S4 and S5**. The nominal resolution was determined based on a Fourier shell correlation cutoff of 0.143.

#### Model building and refinement

Crystal structures of Gβγ and PLCβ3 (PDB IDs 1GP2 and 4GNK, respectively (5, 60)) were rigid-body fit into the cryo-EM map using Chimera (61). The model was then refined using molecular dynamic flexible fitting (MDFF)(62). MDFF configuration files were generated using VMD. During MDFF simulation, Gβγ was set as rigid with domain restraints. The MDFF simulation was conducted with a grid scaling value of 0.5 for 100 ps, followed by 3,000 steps of energy minimization until convergence of the protein RMSD. The MDFF generated model was inspected and manually adjusted in COOT (63), guided by the use of deep-learning-based amino-acid-wise model quality (DAQ) scoring (64, 65) and refined in PHENIX (66). Resulting models were assessed in PHENIX for stereochemical correctness. Maps, half maps, and coordinate files were deposited in the PDB as 9Y7H, 9YAO, and 9YAP and in the EMDB as EMD-72655, EMD-72732, and EMD-72733.

### Activity assays

#### Inositol phosphate accumulation

COS-7 cells were seeded in 12-well culture dishes at a density of 100,000 cells per well and maintained in Dulbecco’s modified Eagle’s medium containing 10% fetal bovine serum (Atlanta Bio), 1X Glutamax (Gibco), 100 units/mL penicillin, and 100 μg/mL streptomycin (Corning) at 37 °C and 5% CO_2_. Cells were transfected with 400 ng of PLCβ3 variant and 200 ng G protein subunit using Fugene 6 (Promega) at a 3:1 ratio per manufacturer’s protocol. Total DNA varied from 700-900 ng per well, with pCMV used as an empty vector. 18-24 h after transfection, the media was changed to low-inositol Ham’s F-10 medium (Gibco) containing 1.5 μCi/well myo-[2-^3^H(N)] inositol (Perkin Elmer) for 16-18 h, then treated with 10 mM LiCl for 1 h to inhibit inositol phosphatases. Media was aspirated, cells were washed twice with PBS, and then lysed by addition of ice-cold 50 mM formic acid. Lysates containing [^3^H]-inositol phosphates were applied to Dowex AGX8 columns to isolate the IP species. Columns were washed twice with 10 CVs of 50 mM formic acid, then 100 mM formic acid, and eluted with three CVs of 1.2 M ammonium formate into scintillation vials containing scintillation fluid and counted.

#### Western blotting

Cells were lysed in SDS sample buffer (100 mM Tris pH 6.8, 6% sucrose, 2% SDS, 715 mM β-mercaptoethanol, and 0.02% bromophenol blue), boiled, and run on a 10% (w/v) SDS–polyacrylamide gel. Proteins were transferred to PDVF for 16 hours at 25 V, followed by incubation with an antibody against PLCβ3 (Cell Signaling Cat: D9D6S) (1:1000), Gβ1 (Thermo: Cat: PA530046) or actin (Cell Signaling: 8H10D10) (1:2000). Goat anti-rabbit HRP or goat anti-rabbit AlexaFlour 800 antibodies (1:10,000) were added before visualizing with ECL reagent (Pierce) for HRP linked antibodies. Western blots were imaged with a GeneGnome imaging system or Azure Sapphire FL respectively.

#### Liposome-based activity assay

Hen egg white phosphatidylethanolamine (PE) at 100 μM and soy phosphatidylinositol (PI) at 250 μM (Avanti Polar Lipids) were resuspended in CHCl_3_, mixed, and dried in 312 μL aliquots in borosilicate glass tubes under N_2_, sealed and stored at −20 °C until use. To prepare liposomes, the lipids were resuspended in 312 μL of sonication buffer (50 mM HEPES pH 7, 80 mM KCl, 2 mM EGTA, and 1 mM DTT) and incubated at room temperature for 5 min, then sonicated to clarity in 30 second duty cycles using a bath sonicator (Avanti Polar Lipids). Each reaction mixture contained 10 μL of liposome solution,10 μL of PLC solution (50 mM HEPES pH 7, 3 mM EGTA, 80 mM KCl, 3 mM DTT, and 3 mg/mL BSA), 5 μL of Gβγ solution or buffer (50 mM HEPES pH 7, 100 mM NaCl, 5 mM MgCl, 1 mM DTT, and 3 mM EGTA), and 5 μL of CaCl_2_ solution (50 mM HEPES pH 7, 3 mM EGTA, 80 mM KCl, 1 mM DTT, and 18 mM CaCl_2_). CaCl_2_ solution was added last to initiate PI hydrolysis upon transfer to 30°C, and incubated for 15 minutes. Control reactions contained all components except CaCl_2_. Reactions were terminated by the addition of an ice-cold quench solution (50 mM HEPES pH 7, 80 mM KCl, 210 mM EGTA, and 1 mM DTT), and incubated on ice.

Inositol phosphate (IP) was quantified using a modified version of the CisBio IP-One Gq assay kit. Following termination, 14 μL of each reaction mixture, 3 μL of d2-labeled IP1, and 3 μL of the cryptate-labeled anti-IP1 antibody (CisBio) were added to a 384-well low-volume white microplate at room temperature (Corning, Corning, NY). Positive controls contained assay buffer, d2-labeled IP1, and cryptate-labeled anti-IP1, and negative controls contained assay buffer, lysis and detection buffers, and cryptate-labeled anti-IP1. The plate was centrifuged at 1000 x *g* for 1 min, incubated at room temperature for 1 h, and fluorescence read on a SynergyNeo2 plate reader (BioTek). The concentration of IP1 was calculated from a standard curve and normalized following the manufacturer’s protocol (CisBio).

### BRET

For Venus-Gβγ interaction experiments HEK 293 cells were transfected with 0.01 μg HiBit-PLCβ, 0.1 μg CMV-LgBit, 0.4 μg Gα_q_, 0.3 μg Venus-1-155-Gγ2, 0.3 μg Venus-155-239-Gβ1 and 0.3 μg SNAPf-AGTR1. For Gα_q_-Venus interaction experiments, Gα_q_-Venus replaced Gα_q_, and unlabeled Gβ1 and Gγ2 replaced their Venus-labeled counterparts. For PI(4,5)P2 hydrolysis assays, HEK 293 cells were transfected with 0.01 μg Nluc-PH, 0.1 μg SNAPf-AGTR1 and 0.8 μg of mem-link-Venus. For PI(4,5)P2 hydrolysis reconstitution assays, ΔPLC cells were transfected with the same components plus 0.01 μg of HiBit-PLCβ, HiBit-PLCβ-CAAX or variants thereof. To sequester either active Gα_q_ or Gβγ, GRK2RH or GRK3ct were added at 0.2 μg per well, respectively. For translocation bystander BRET experiments, HEK 293 cells were transfected with 1 μg mem-link-Venus, 0.1 μg CMV-LgBiT, 0.3 μg SNAPf-AGTR1, 0.4 Gα_q_ μg and 0.01 μg HiBit-PLCβ or variant. For all experiments, cells were washed and resuspended in Dulbecco’s phosphate buffered saline (DPBS) and distributed to 96-well plates in suspension immediately before taking BRET measurements. All BRET measurements were made in the presence of furimazine (Promega or ChemShuttle; 1:1,000 as supplied or from a 5 mM stock dissolved in 90% ethanol/10% glycerol). BRET and luminescence measurements were made using a Polarstar Optima or Lumistar Omega plate reader (BMG Labtech); angiotensin II was injected from a 10-fold concentrated solution. Raw BRET signals were calculated as the emission intensity at 520–545 nm divided by the emission intensity at 475–495 nm. Net BRET signals were calculated as the raw BRET signal minus the raw BRET signal measured from cells expressing only the donor.

### Statistical analysis

All analysis was carried out using GraphPad Prism Version 10.5.0.

## Materials and data availability

All study data are included in the article and/or supporting information. All unique materials produced for this study are available from the corresponding author upon request.

## Acknowledgements and funding sources

We thank Steve Wilson and Drs. John. J. G. Tesmer, Val J. Watts, Thomas Klose, and Frank Vago for technical and conceptual assistance. A.I. was funded by Japan Society for the Promotion of Science (JP24K21281 and JP25H01016); Japan Science and Technology Agency (JPMJFR215T and JPMJMS2023); Japan Agency for Medical Research and Development (JP22ama121038 and JP22zf0127007); and The Uehara Memorial Foundation. E.K. was supported by the Deutsche Forschungsgemeinschaft (DFG, German Research Foundation) with the grant 214362475/GRK1873/3. This work is supported by F32GM145110-01 to I.J.F, R35GM145284 to N.A.L., 1R01HL141076 and 1R01GM152701 to A.M.L.

## Supplemental Information

**Figure S1.**
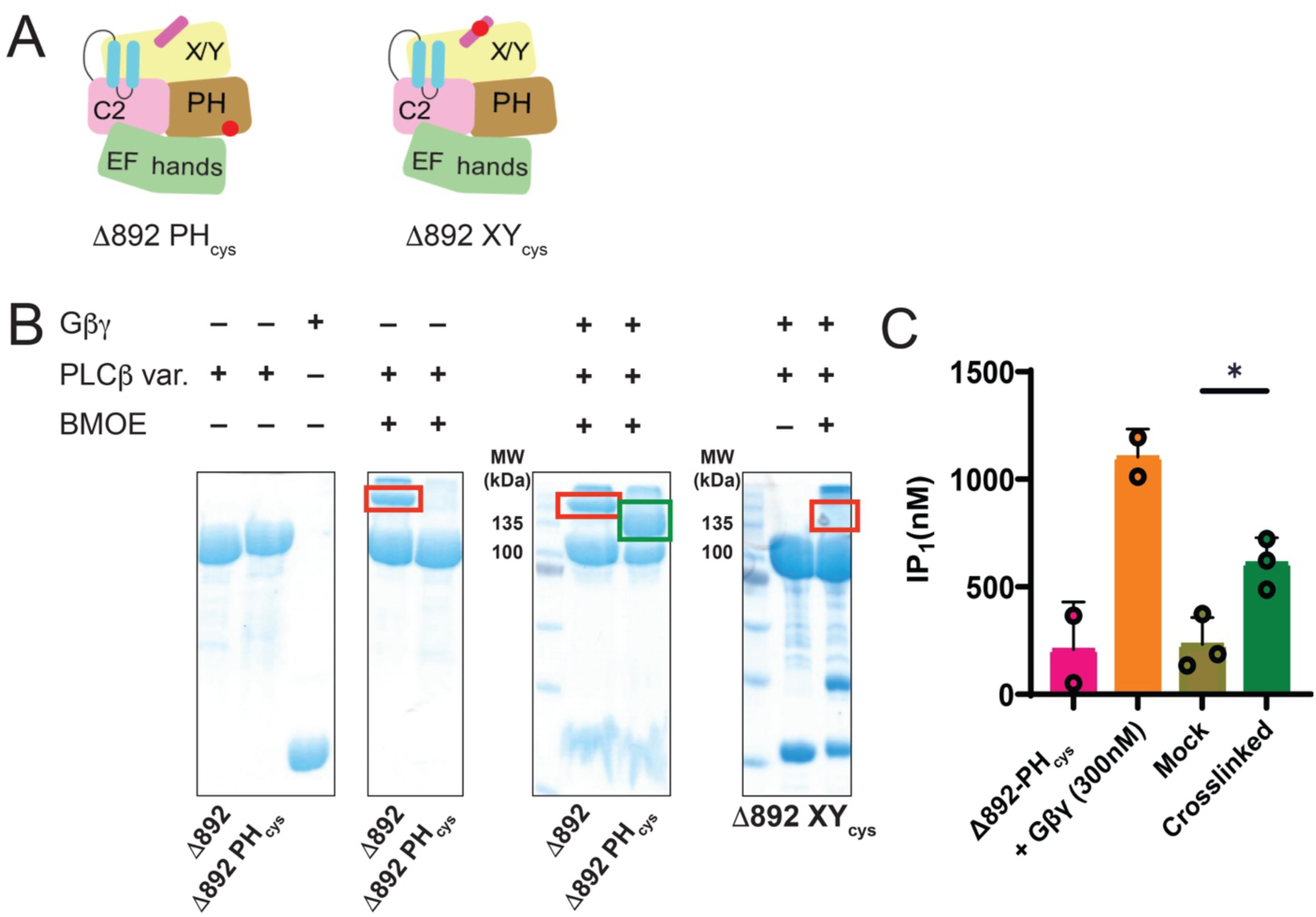
Isolation of a Crosslinked and Functional Gβγ–PLCβ3 complex. (**A**) Schematic of PLCβ3 Δ892 variants used for crosslinking studies. (**B**) Representative crosslinking experiments with Gβγ C68S and PLCβ3-Δ892 variants analyzed by SDS-PAGE and stained with Coomassie. PLCβ3-Δ892 undergoes extensive self-crosslinking (red box, left). Mutation of solvent-exposed cysteines and introduction of a single cysteine in the PH domain (E60C) eliminates self-crosslinking and allows crosslinking between Gβγ C68S and PLCβ3-Δ892_PH_ (green box). PLCβ3-Δ892 XY_cys_, which lacks solvent-exposed cysteines except for C516 in the flexible X–Y linker, eliminated self-crosslinking but failed to crosslink to Gβγ-C68S. (**C**) BMOE-crosslinked complexes between wild-type Gβγ and PLCβ3 Δ892_PH_ have higher activity in a liposome-based assay than the uncrosslinked control. Data are the mean of three separate experiments ± SD. Mock crosslinked and crosslinked samples were compared using an unpaired T-test. *, p < 0.05.

**Figure S2.**
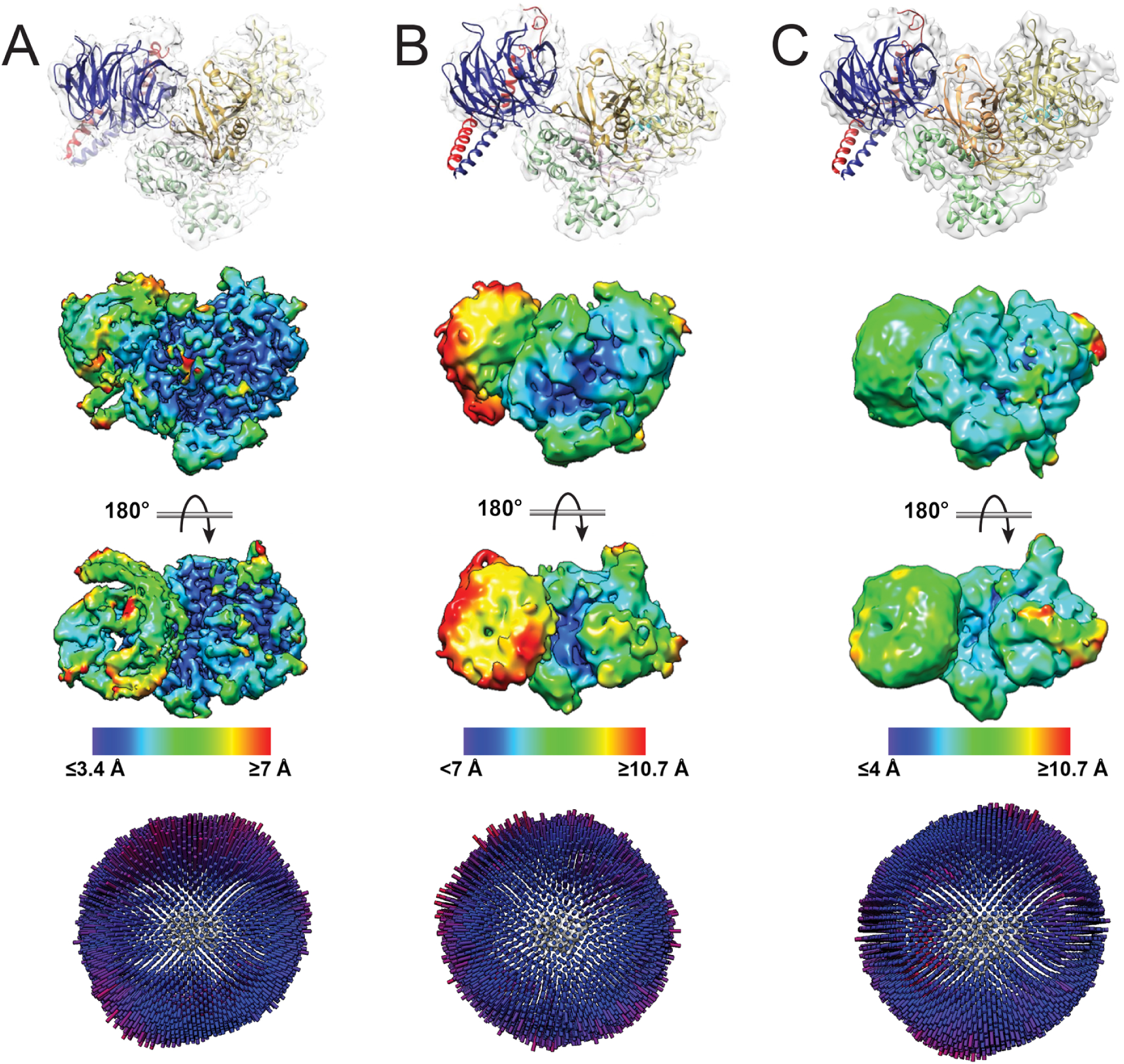
Cryo-EM densities of Gβγ–PLCβ3 Δ892-PH_cys_ complexes. (**A**) *Top.* Model of the larger particle population in the BMOE-crosslinked Gβγ–PLCβ3 Δ892-PH_cys_ reconstruction fit into the cryo-EM density map. *Bottom*. Cryo-EM map colored by local resolution. (**B**) Model of the smaller particle population in the BMOE-crosslinked Gβγ–PLCβ3 Δ892-PH_cys_ reconstruction fit into the cryo-EM density map. *Bottom.* Cryo-EM map colored by local resolution. (**C**) *Top.* Model of the BM(PEG)2-crosslinked Gβγ–PLCβ3 Δ892-PH_cys_ reconstruction fit into the cryo-EM density map. *Bottom*. Cryo-EM map colored by local resolution. In all reconstructions, resolution is lower for Gβγ, consistent with a dynamic interface in solution.

**Figure S3.**
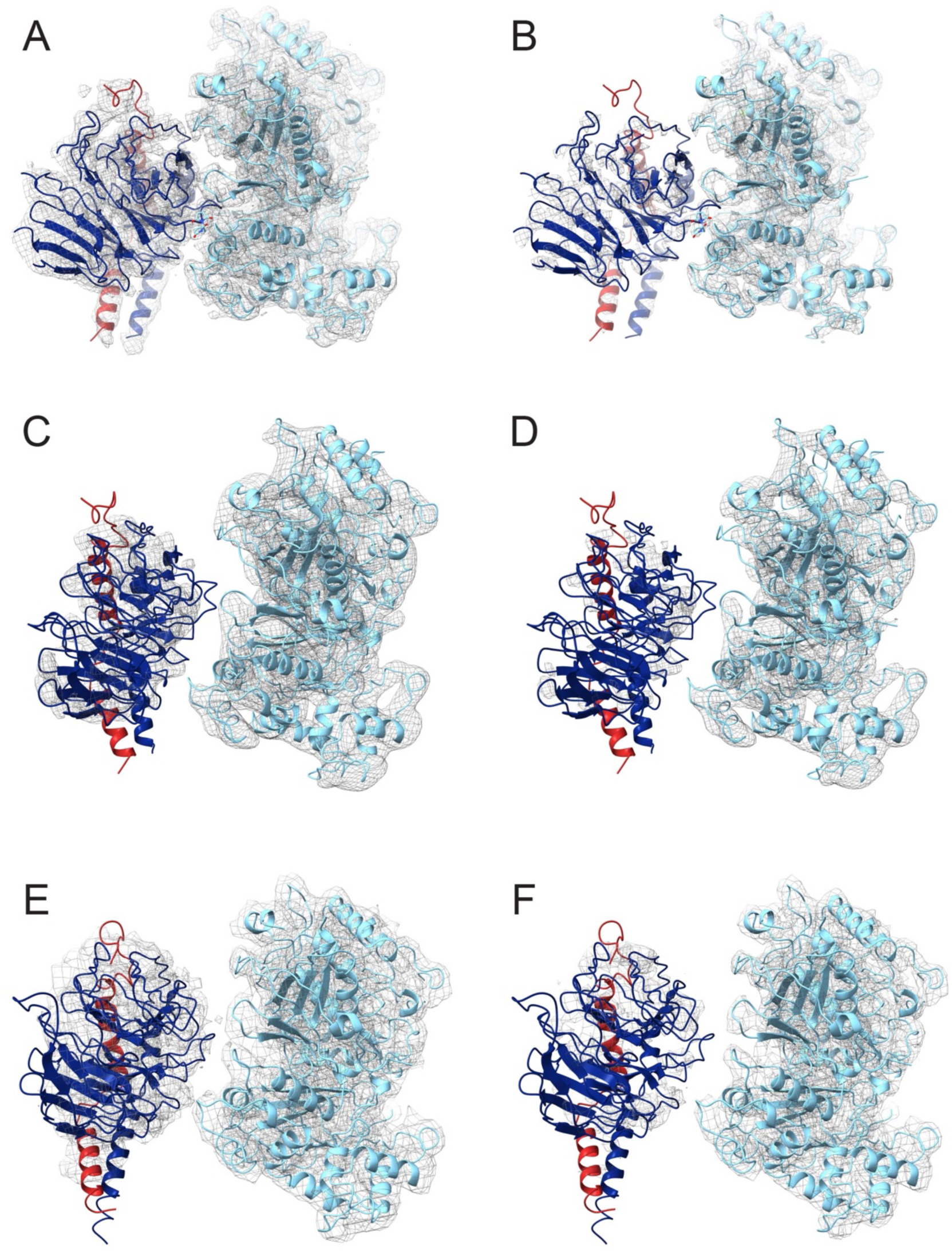
Contoured cryo-EM density maps of the Gβγ–PLCβ3 Δ892-PH_cys_ complexes. The 4 Å Gβγ–PLCβ3 Δ892-PH_cys_ complex fit in the density map contoured to (**A**) 0.197 σ or (**B**) 0.297 σ. The 7 Å Gβγ–PLCβ3 Δ892-PH_cys_ complex contoured to (**C**) 0.197σ or (**D**) 0.227 σ, and the 4.5 Å BM-PEG crosslinked Gβγ–PLCβ3-Δ892 PH_cys_ complex fit in density contoured to (**D**) 0.040 σ or (**E**) 0.050 σ.

**Figure S4.**
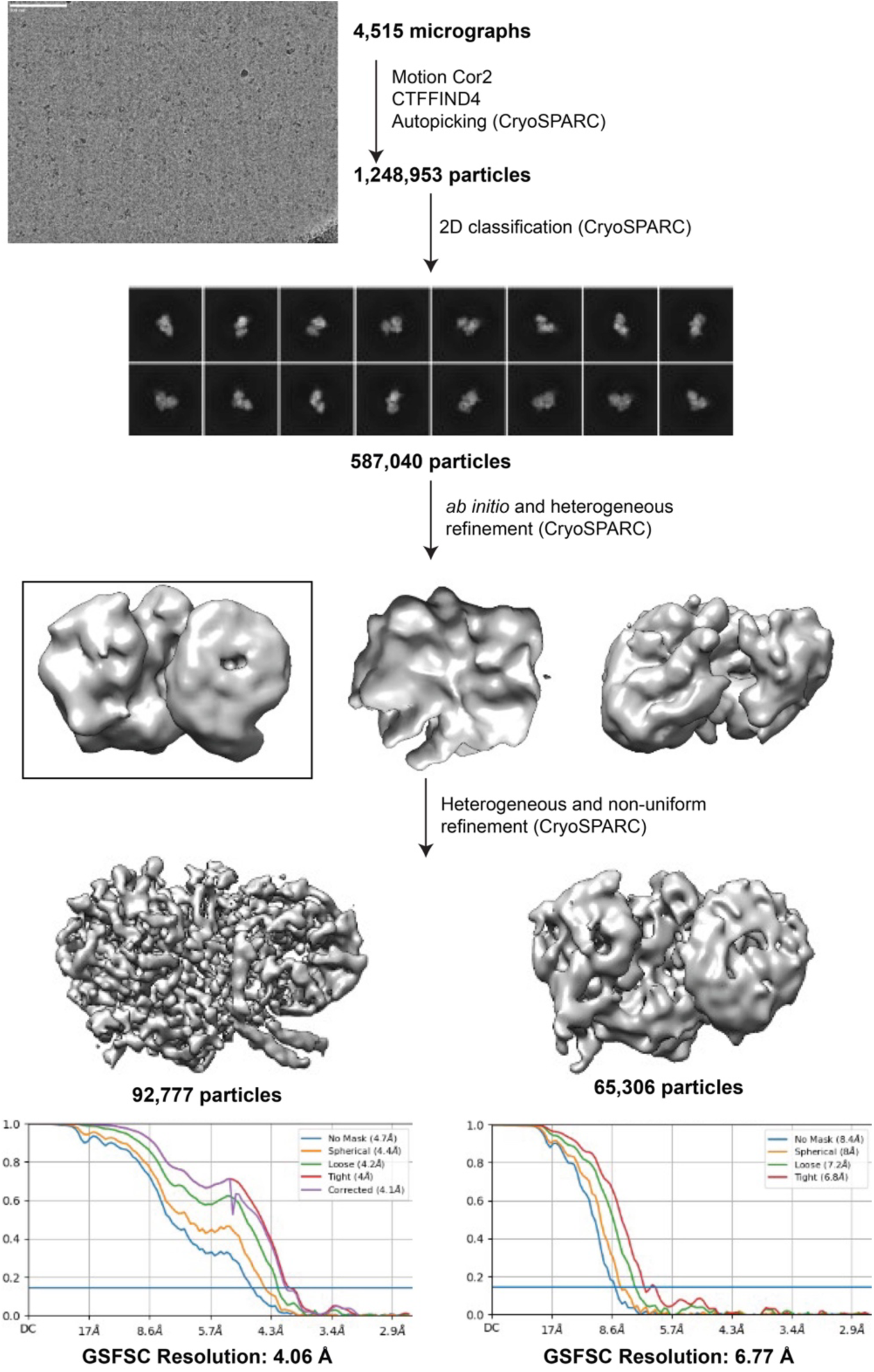
Cryo-EM data workflow and resolution analysis of the BMOE-crosslinked Gβγ–PLCβ3 Δ892-PH_cys_ complex. The workflow, including a representative micrograph, 2D class averages (box size: 276 Å) and Fourier shell correlation (FSC) curves calculated from two independent reconstructions by CryoSPARC(58). The nominal resolution of the resulting map, as defined by the 0.143 cutoff, is indicated by the horizontal blue line.

**Figure S5.**
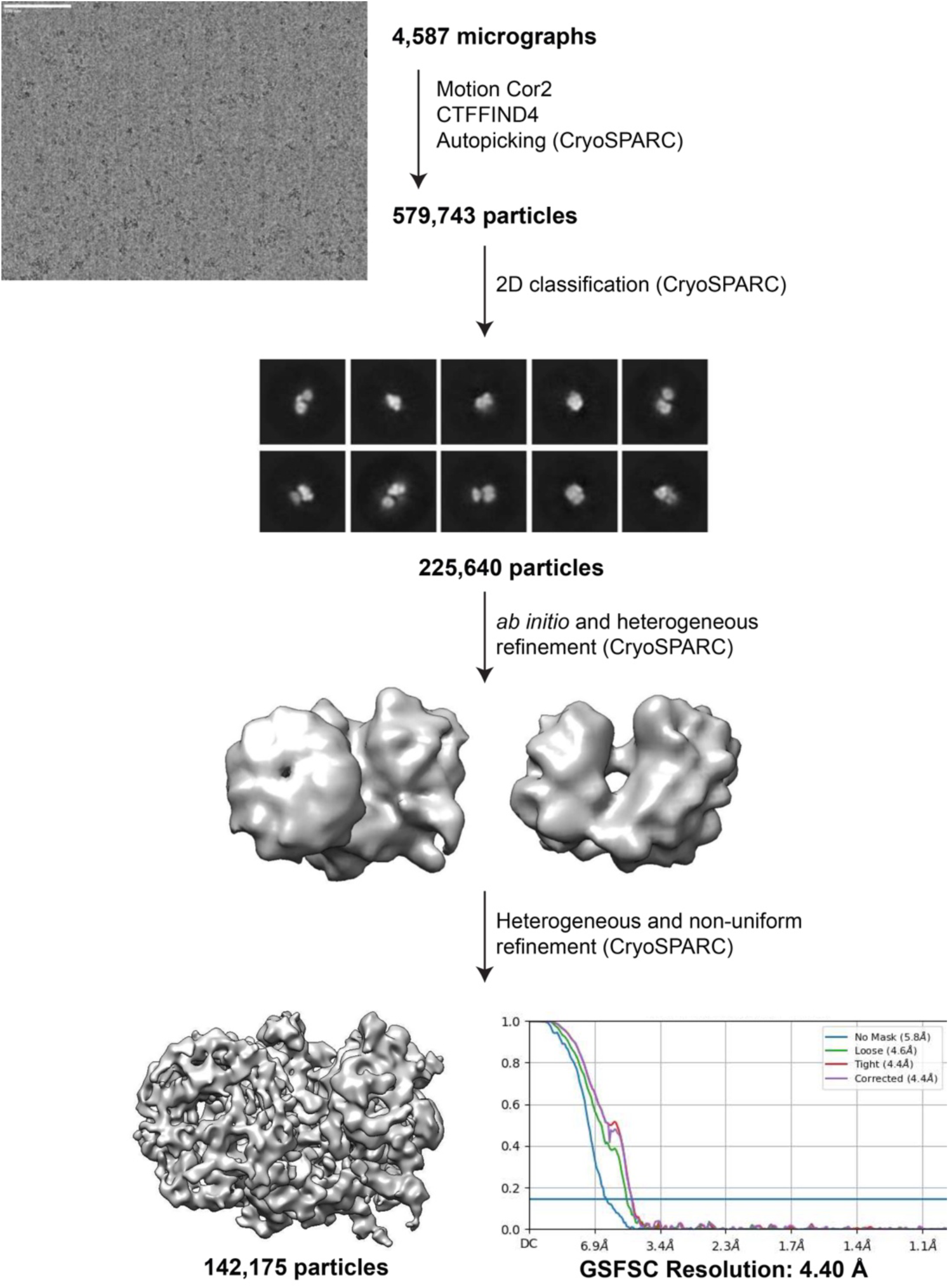
Cryo-EM data workflow and resolution analysis of the BMPEG-crosslinked Gβγ–PLCβ3 Δ892-PH_cys_ complex. The workflow, including a representative micrograph, 2D class averages (box size: 276 Å) and Fourier shell correlation (FSC) curves calculated from two independent reconstructions by CryoSPARC(58). The nominal resolution of the resulting map, as defined by the 0.143 cutoff, is indicated by the horizontal blue line.

**Figure S6.**
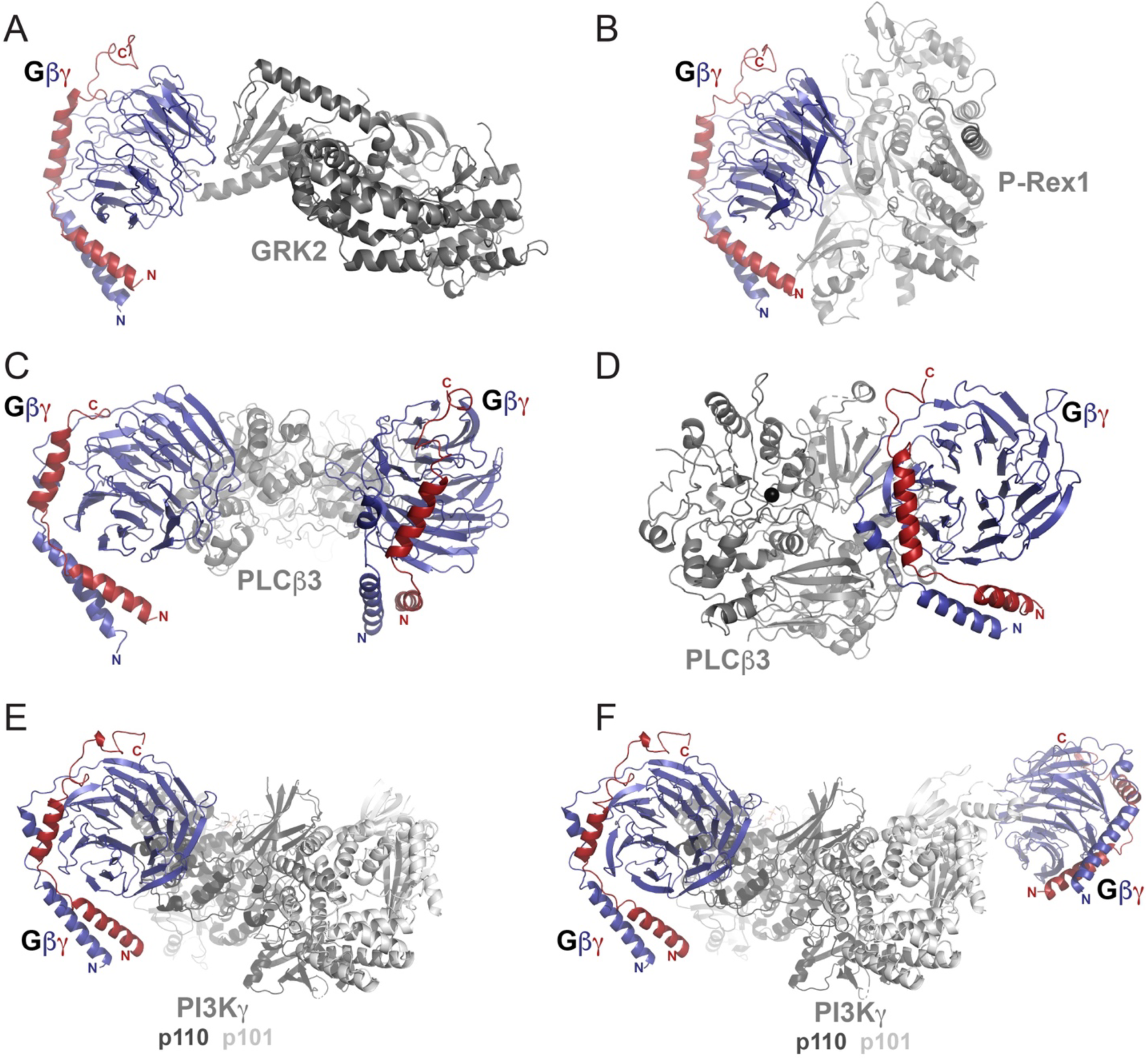
Gβγ–effector enzyme complexes. Experimentally determined structures of Gβγ in complex with effector enzymes including (A) GPCR kinase 2 (GRK2, PDB ID 3CIK), (B) phosphatidylinositol-3,4,5-trisphosphate dependent Rac exchange factor 1 (P-Rex 1, PDB ID 6PCV), (C) liposome-tethered PLCβ3 (PDB ID 8EMW), (D) PLCβ3 in solution (this work, PDB ID 9Y7H), (E) the high affinity site of phosphatidylinositol 3-kinase γ (PI3Kγ, PDB ID 8SOC), and (F) both sites of PI3Kγ (PDB ID 8SOD). In all structures, Gβ is shown in dark blue, Gγ in navy, and the effector enzyme in gray. In the Gβγ–PI3K–Gβγ structures, the p110 subunit is shown in dark gray, and the p101 subunit in light gray.

**Figure S7.**
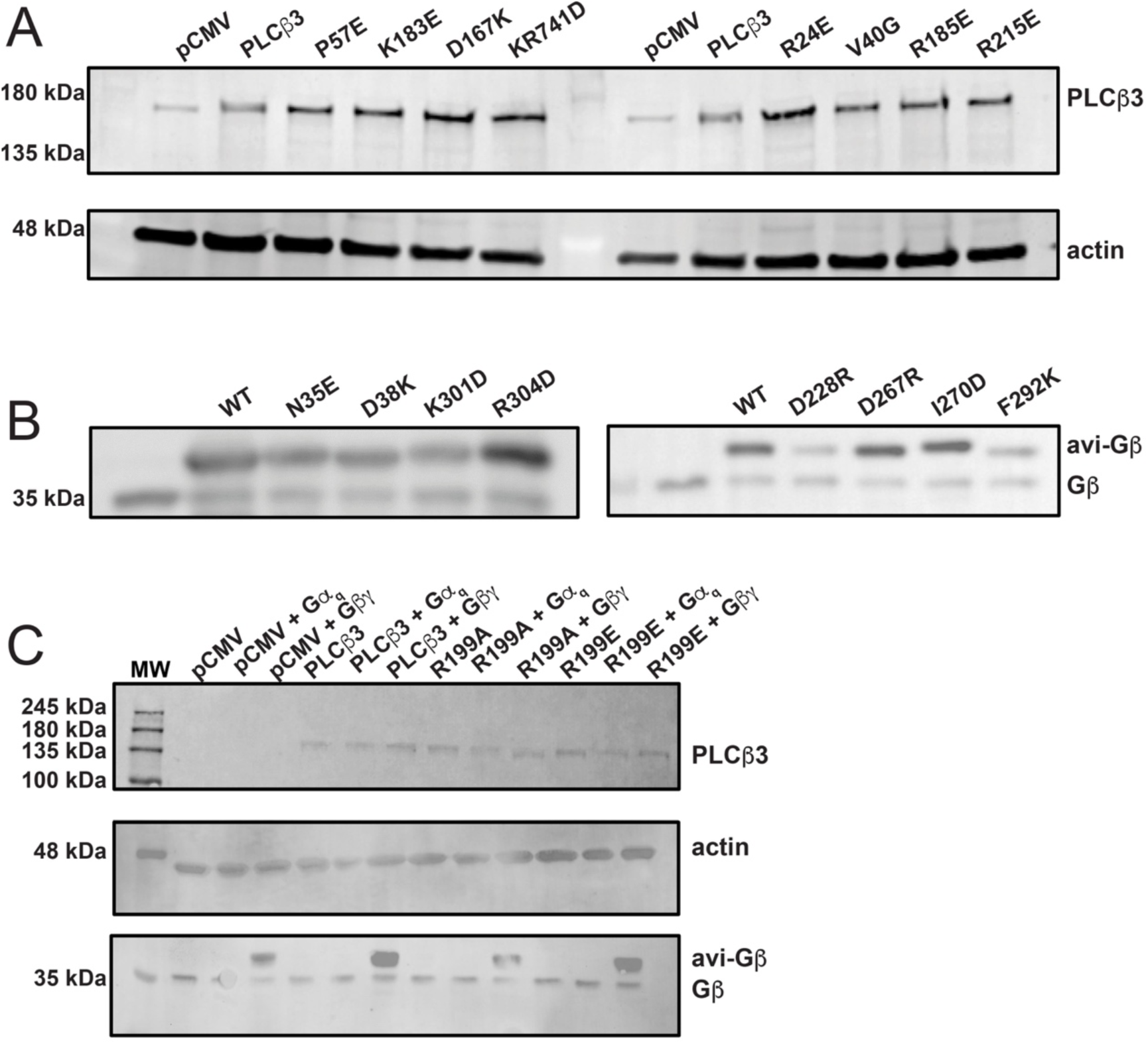
Expression of PLCβ3 and Gβγ mutants. Representative western blot image of cell lysates containing mutants of interest. (**A**) Western blot image of PLCβ3 mutants. (**B**) Western blot image of Gβ1 mutants. Transfected Gβγ is Avi-tagged (avi-Gβγ) and endogenous Gβ is detected in the untransfected controls. (**C**) Representative western blot image of cell lysates containing PLCβ3 and the R199A and R199E mutants (top) and co-transfected avi-Gβ and endogenous Gβ (below)

**Figure S8.**
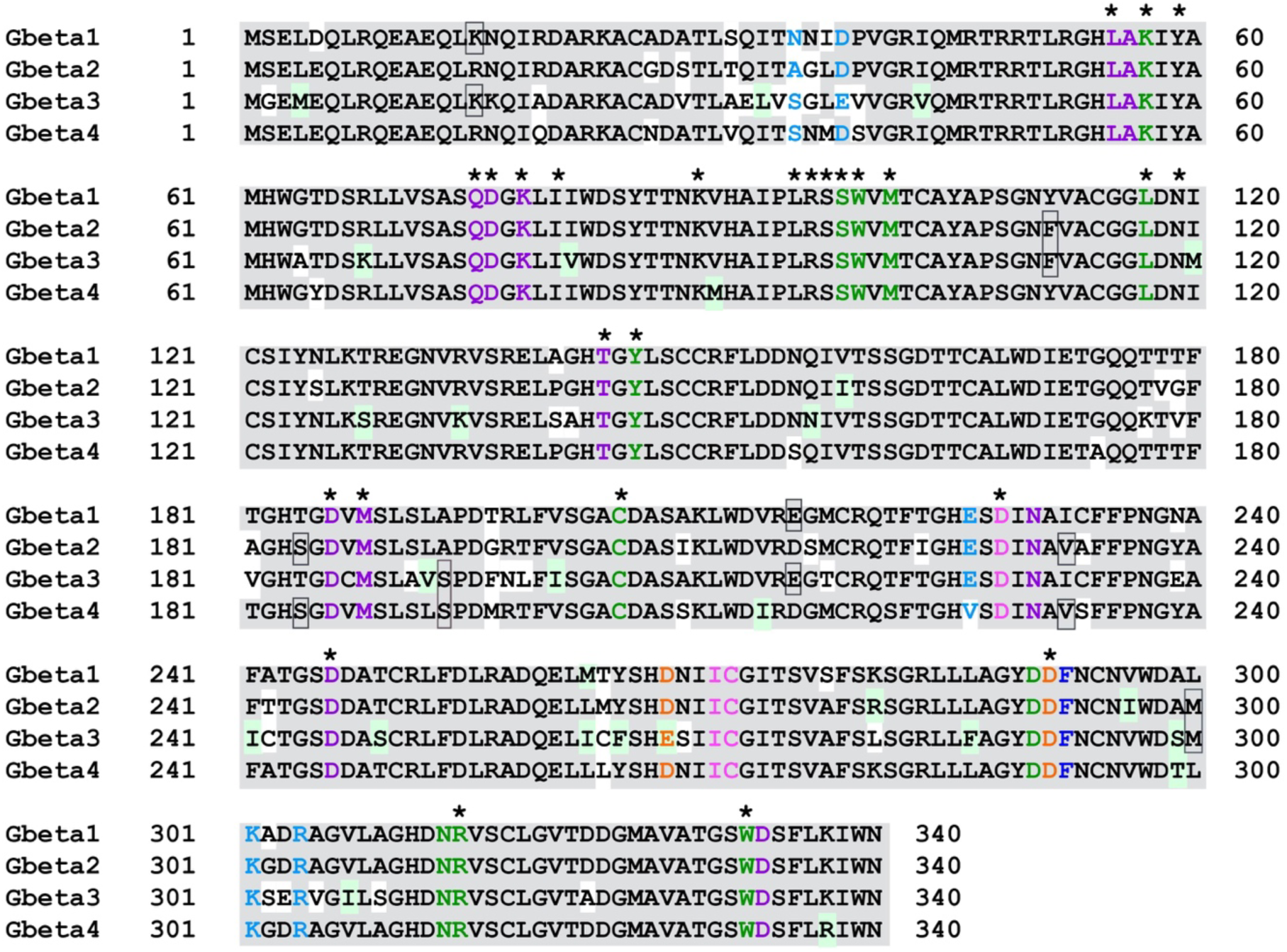
PLCβ binding surfaces are conserved in human Gβ1-4 isoforms. Sequence alignment of human Gβ1-4 subunits. Residues highlighted in light gray are conserved in all isoforms and those highlighted in light green are share similar properties. Residues in orange and purple are those that bind only to the PLCβ3 PH domain or to EF hands, respectively, in the liposome-tethered reconstruction. Residues in green interact with PLCβ3 in both sites. Residues in cyan are unique to the crosslinked complex described in this work. Residues in dark blue interact with the PLCβ3 PH domain in the crosslinked and liposome-tethered reconstructions. Residues in hot pink are common to all three observed Gβγ-PLCβ3 interfaces. Asterisks correspond to residues known participate in other Gβγ–effector enzyme interactions. Sequences correspond to UNIPROT entries P62874 (Gβ1), P62879 (Gβ2), P16520 (Gβ3), and Q9HAV0 (Gβ4).

**Figure S9.**
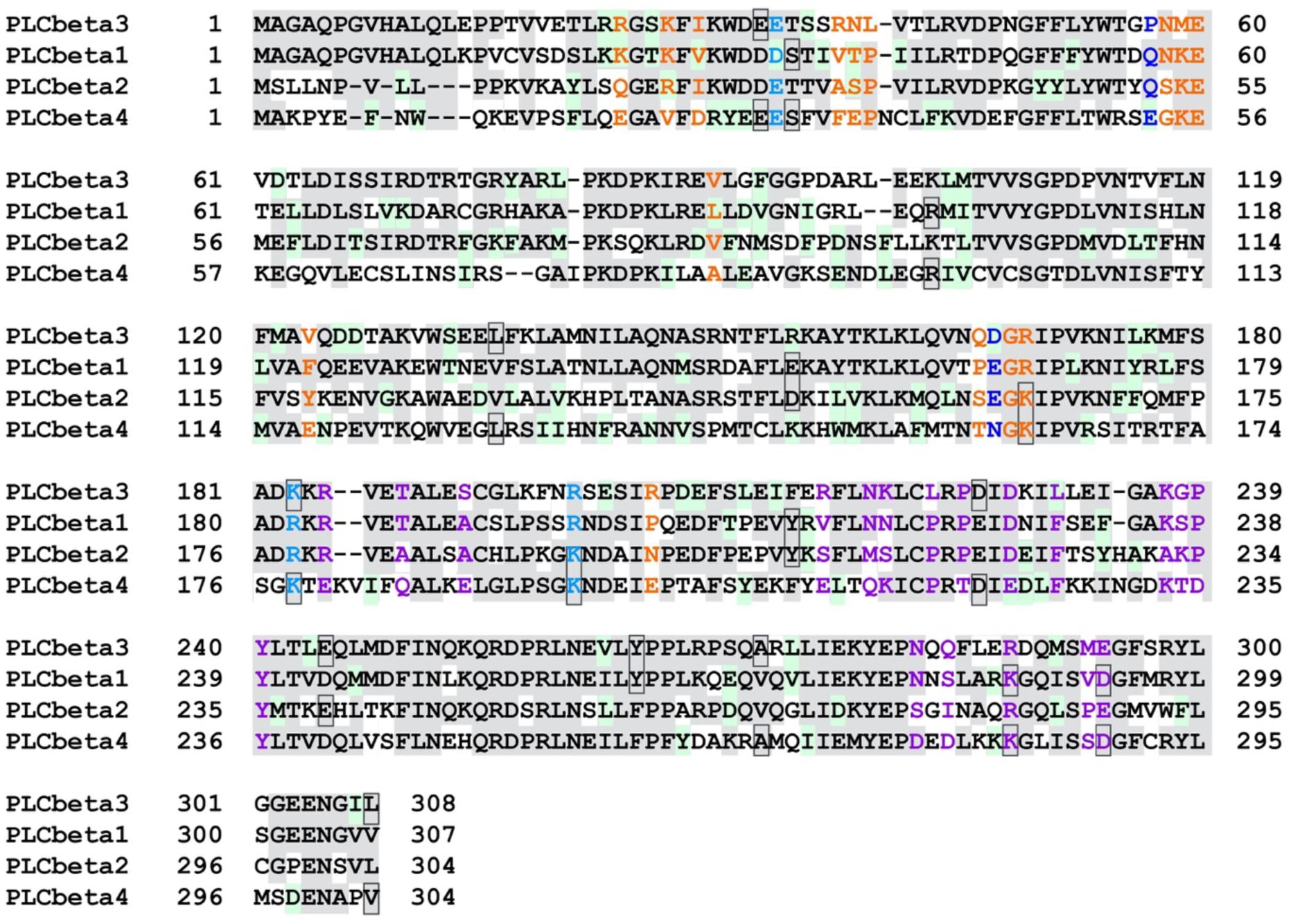
Gβγ binding residues are not well conserved across the human PLCβ isoforms. Sequence alignment of the human PLCβ PH and EF hand domains. Conserved residues are highlighted in light gray and residues with similar properties are highlighted in light green. Residues in the PLCβ3 PH domain that bind Gβγ are in orange, and those that bind Gβγ through the EF hands are in purple. PLCβ3 residues that interact with Gβγ in the crosslinked structure are in cyan. Residues that interact with Gβγ in both cryo-EM structures are in dark blue. Sequences correspond to UNIPROT entries Q9NQ66 (PLCβ1), Q00722 (PLCβ2), Q01970 (PLCβ3), and Q15147 (PLCβ4).

**Figure S10.**
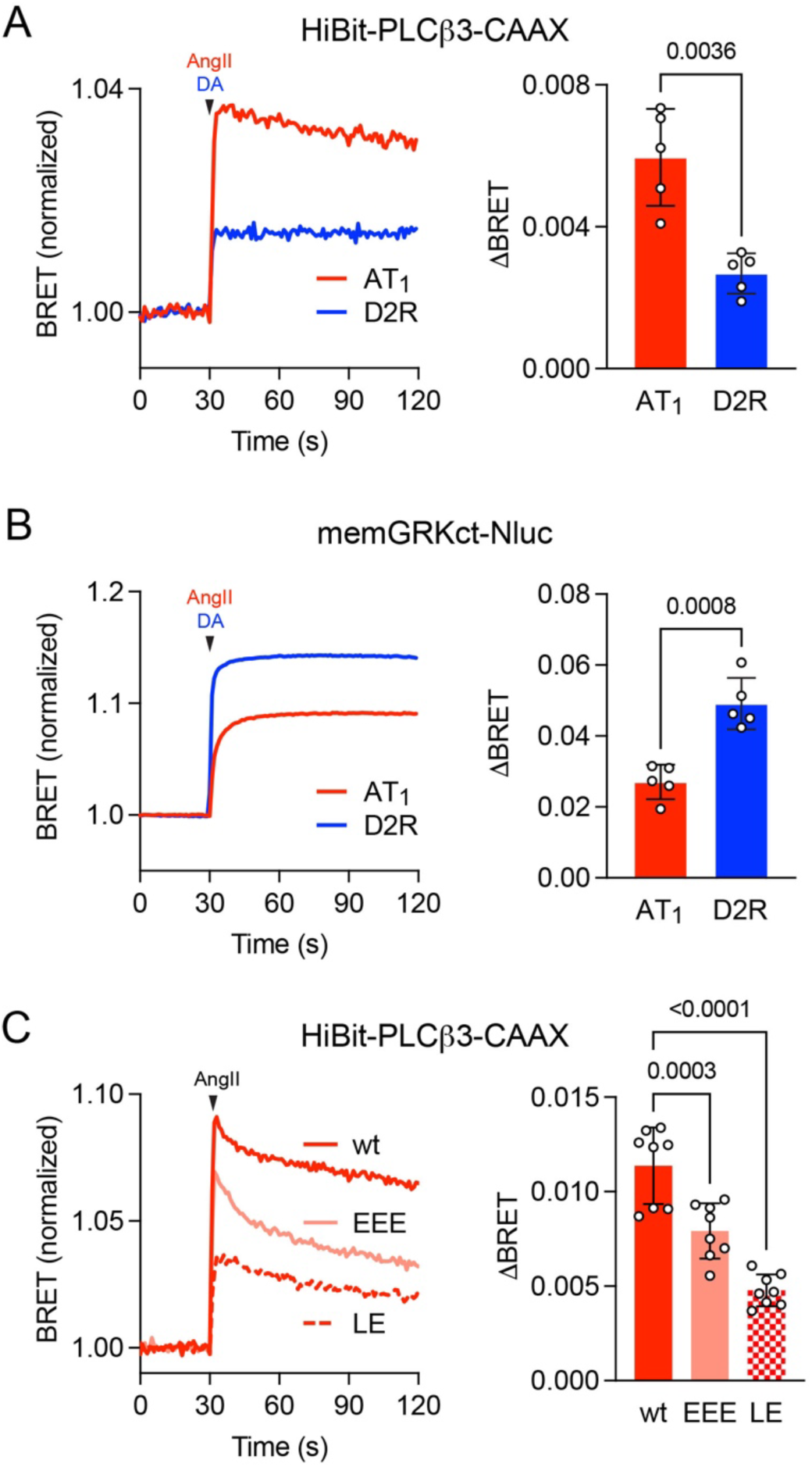
Gα_q_ binding promotes Gβγ binding to PLCβ3. (**A**) BRET between membrane-anchored HiBit-PLCβ3-CAAX and Venus-Gβγ increases after activation of AT_1_ with angiotensin II (AngII; 1 μM) or dopamine D2R receptors with dopamine (DA; 100 μM). Traces represent the average of twenty replicates from five independent experiments. Signals were significantly smaller (p=0.0036) after activation of D2R; Welch’s t-test. (**B**) BRET between the Gβγ sensor memGRKct-Nluc and Venus-Gβγ increases after activation of AT_1_ or D2R. Traces represent the average of twenty replicates from five independent experiments. Signals were significantly larger (p=0.0008) after activation of D2R; Welch’s t-test. Transfection was identical for panels A and B with the substitution of memGRKct-Nluc for HiBit-PLCβ3-CAAX in panel B; Gα_q_ and Gα_i1_ were overexpressed together with Venus-Gβγ in both panels. (**C**) BRET between HiBit-PLCβ3-CAAX wild-type (wt) and mutants with defective Gα_q_ binding to the distal CTD (EEE) or proximal CTD (LE). Traces represent the average of 32 replicates from eight independent experiments. Signals were significantly smaller for both mutants; one-way ANOVA with Dunnett’s post-hoc comparisons.

**Figure S11.**
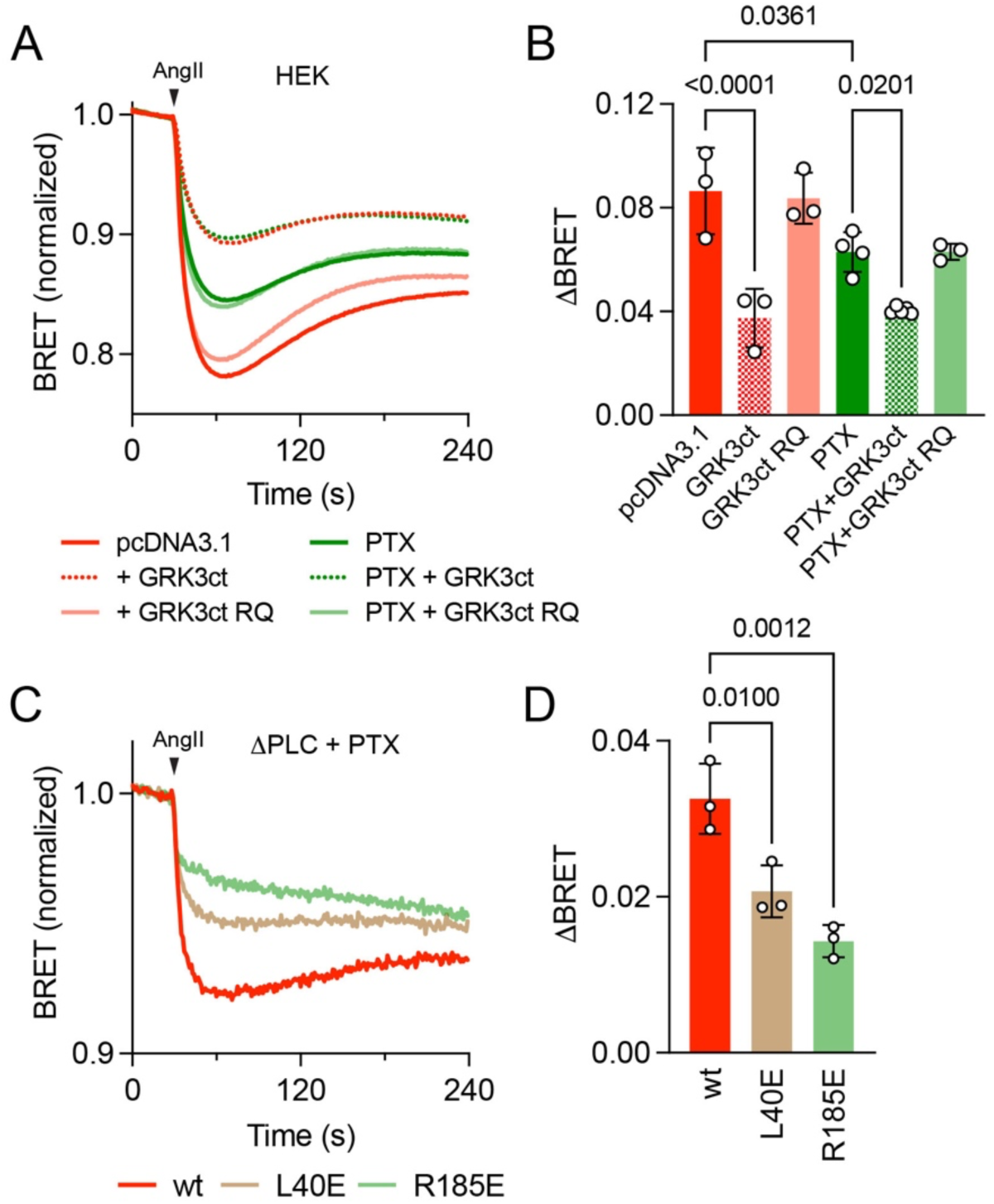
PLCβ-mediated PI(4,5)P2 hydrolysis is facilitated by Gβγ derived from both G_q_ and G_i/o_ heterotrimers. (**A**) In HEK cells, bystander BRET between Nluc-PH and mem-Venus decreases in response to AT1 activation with AngII (1 μM). Responses are inhibited by membrane-tetheredGRK3ct, which sequesters Gβγ, but not the binding-defective R587Q mutant (GRKct RQ), both before and after inactivation of G_i/o_ heterotrimers with pertussis toxin (PTX). Traces are the average twelve-sixteen replicates from 3-4 independent experiments. (**B**) Grouped data from the same experiments as panel A; indicated p values are from one-way ANOVA with Tukey’s multiple comparisons test. (**C**) In ΔPLC cells expressing PTX, HiBit-PLCβ3 L40E and R185E mutants fail to fully reconstitute AngII-induced PI(4,5)P2 hydrolysis compared to the wild-type (wt) enzyme; traces are the average of twelve replicates from three independent experiments. (**D**) Grouped data from the same experiments as panel C; indicated p values are from one-way ANOVA with Dunnett’s multiple comparisons test.

**Table S1.**
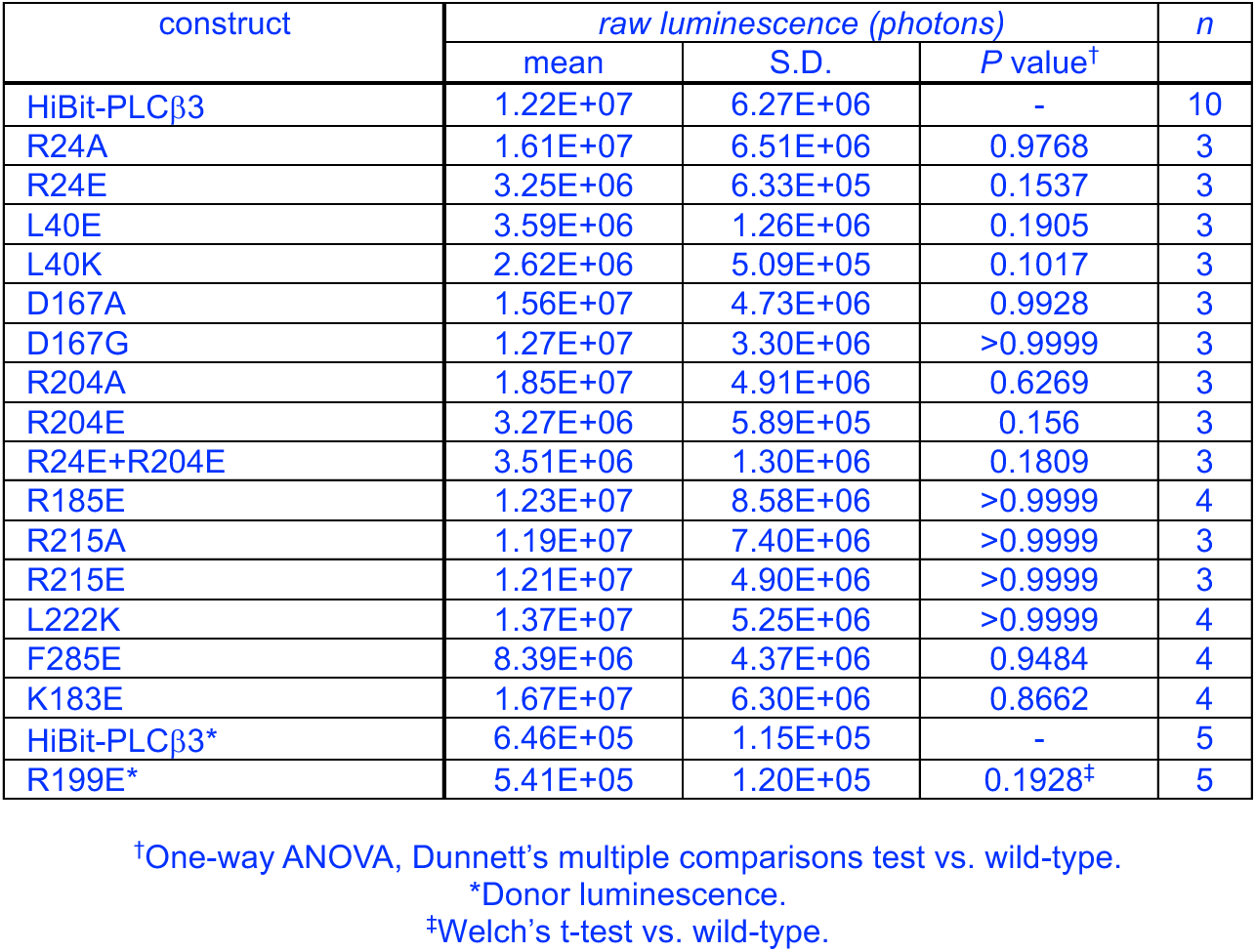
Expression of HiBit-PLCβ3 variants as indicated by LgBit-complemented luminescence in intact cells.

**Table S2.**
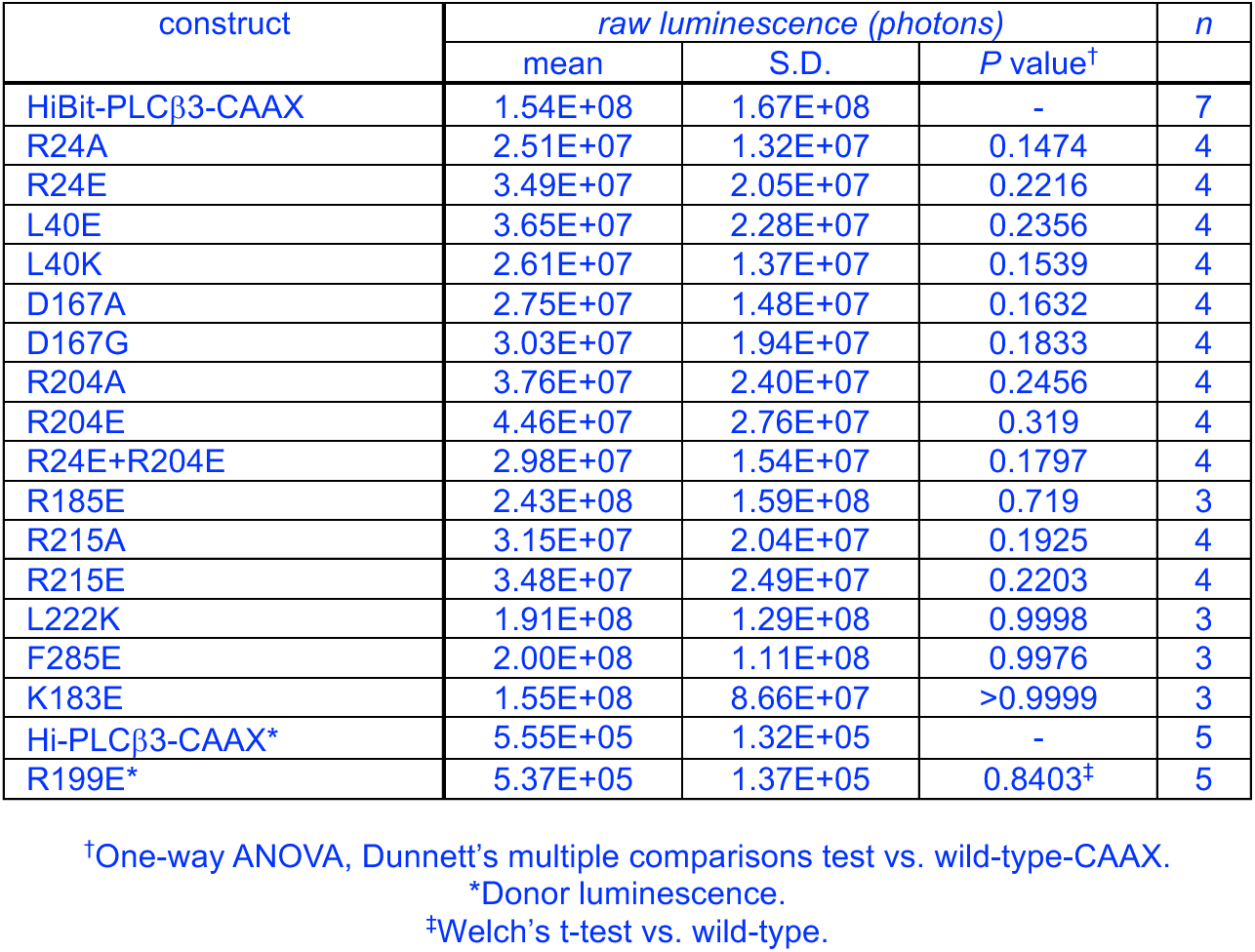
Expression of HiBit-PLCβ3-CAAX variants as indicated by LgBit-complemented luminescence in intact cells.

**Table S3.** Angiotensin II-induced BRET between HiBit-PLCβ3-CAAX variants and Venus-Gβγ.

| construct | $\Delta BRET$ | | | *n* |
| --- | --- | --- | --- | --- |
|  | mean | S.D. | *P* value<sup>†</sup> |  |
| HiBit-PLCβ3-CAAX | 0.01163 | 0.004365 | - | 10 |
| +GRK3ct | -0.000968 | 0.001502 | <0.0001 | 8 |
| R24A | 0.008984 | 0.001371 | 0.9249 | 5 |
| R24E | 0.004995 | 0.00116 | 0.0040 | 7 |
| L40E | 0.001863 | 0.00054 | <0.0001 | 5 |
| L40K | 0.002092 | 0.00071 | <0.0001 | 5 |
| D167A | 0.0306 | 0.01035 | <0.0001 | 5 |
| D167G | 0.01943 | 0.004322 | 0.0023 | 5 |
| R204A | 0.007136 | 0.001778 | 0.2798 | 5 |
| R204E | 0.003198 | 0.000906 | <0.0001 | 7 |
| R24E+R204E | 0.002536 | 0.001165 | 0.0008 | 4 |
| R185E | 0.000937 | 0.000516 | 0.0004 | 3 |
| R215A | 0.02478 | 0.00525 | <0.0001 | 5 |
| R215E | 0.02706 | 0.006269 | <0.0001 | 5 |
| L222K | 0.006882 | 0.00098 | 0.4898 | 3 |
| F285E | 0.002372 | 0.000248 | 0.0033 | 3 |
| K183E | 0.0116 | 0.002322 | >0.9999 | 4 |
| R199E | 0.003528 | 0.001421 | 0.0041 | 4 |

<sup>†</sup>One-way ANOVA, Dunnett's multiple comparisons test vs. wild-type-CAAX.

**Table S4.** Angiotensin II-induced PI(4,5)P2 hydrolysis mediated by HiBit-PLCβ3 variants.

| construct | $\Delta BRET$ | | | *n* |
| --- | --- | --- | --- | --- |
|  | mean | S.D. | *P* value<sup>†</sup> |  |
| HiBit-PLCβ3 | 0.05237 | 0.005064 | - | 11 |
| +GRK3ct | 0.02145 | 0.003139 | <0.0001 | 11 |
| R24A | 0.053 | 0.008833 | >0.9999 | 3 |
| R24E | 0.04107 | 0.004471 | 0.0008 | 5 |
| L40E | 0.02582 | 0.004777 | <0.0001 | 5 |
| L40K | 0.02727 | 0.003389 | <0.0001 | 5 |
| D167A | 0.07282 | 0.001987 | <0.0001 | 3 |
| D167G | 0.06207 | 0.01019 | 0.0520 | 3 |
| R204A | 0.04634 | 0.002524 | 0.6015 | 3 |
| R204E | 0.03782 | 0.007242 | <0.0001 | 5 |
| R24E+R204E | 0.03578 | 0.0041 | <0.0001 | 4 |
| R185E | 0.01643 | 0.001868 | <0.0001 | 3 |
| R215A | 0.06183 | 0.005277 | 0.0636 | 3 |
| R215E | 0.06154 | 0.006815 | 0.0809 | 3 |
| L222K | 0.02474 | 0.001306 | <0.0001 | 3 |
| F285E | 0.01552 | 0.00182 | <0.0001 | 3 |
| K183E | 0.03727 | 0.009687 | 0.0002 | 3 |
| R199E | 0.02955 | 0.003595 | <0.0001 | 5 |

<sup>†</sup>One-way ANOVA, Dunnett's multiple comparisons test vs. wild-type.

**Table S5.** Angiotensin II-induced BRET between HiBit-PLCβ3 variants and Gα_q_-Venus.

| construct | $\Delta BRET$ | | | *n* |
| --- | --- | --- | --- | --- |
|  | mean | S.D. | *P* value<sup>†</sup> |  |
| HiBit-PLCβ3 | 0.09643 | 0.02322 | - | 3 |
| R24A | 0.08169 | 0.01774 | 0.9344 | 3 |
| R24E | 0.09379 | 0.01895 | >0.9999 | 3 |
| L40E | 0.105 | 0.01828 | 0.9996 | 3 |
| L40K | 0.07575 | 0.008115 | 0.6243 | 3 |
| D167A | 0.1023 | 0.01475 | >0.9999 | 3 |
| D167G | 0.05385 | 0.01087 | 0.0103 | 3 |
| R204A | 0.0951 | 0.02177 | >0.9999 | 3 |
| R204E | 0.1066 | 0.01497 | 0.9973 | 3 |
| R24E+R204E | 0.1008 | 0.01792 | >0.9999 | 3 |
| R185E | 0.1094 | 0.02068 | 0.9753 | 3 |
| R215A | 0.1116 | 0.01287 | 0.9203 | 3 |
| R215E | 0.08856 | 0.01038 | 0.9999 | 3 |
| L222K | 0.08054 | 0.01358 | 0.8935 | 3 |
| F285E | 0.05337 | 0.001885 | 0.0092 | 3 |
| K183E | 0.1021 | 0.007104 | >0.9999 | 3 |
| R199E | 0.1259 | 0.01843 | 0.0620 | 5 |

<sup>†</sup>One-way ANOVA, Dunnett's multiple comparisons test vs. wild-type.

**Table S6.** Angiotensin II-induced BRET between HiBit-PLCβ3 variants and mem-Venus.

| construc | $\Delta BRET$ | | | *n* |
| --- | --- | --- | --- | --- |
|  | mean | S.D. | *P* value<sup>†</sup> |  |
| HiBit-PLCβ3 | 0.03869 | 0.005318 | - | 3 |
| R24A | 0.03012 | 0.00526 | 0.0619 | 3 |
| R24E | 0.03258 | 0.002824 | 0.3757 | 3 |
| L40E | 0.03697 | 0.003716 | >0.9999 | 3 |
| L40K | 0.03456 | 0.002387 | 0.8565 | 3 |
| D167A | 0.03445 | 0.003044 | 0.8352 | 3 |
| D167G | 0.02822 | 0.003124 | 0.0108 | 3 |
| R204A | 0.03021 | 0.006573 | 0.0667 | 3 |
| R204E | 0.03401 | 0.004711 | 0.7333 | 3 |
| R24E+R204E | 0.03131 | 0.003755 | 0.1608 | 3 |
| R185E | 0.02671 | 0.002865 | 0.0023 | 3 |
| R215A | 0.0381 | 0.004039 | >0.9999 | 3 |
| R215E | 0.03365 | 0.005771 | 0.6405 | 3 |
| L222K | 0.03058 | 0.003968 | 0.0911 | 3 |
| F285E | 0.02902 | 0.002389 | 0.0232 | 3 |
| K183E | 0.02648 | 0.002579 | 0.0018 | 3 |
| R199E | 0.03633 | 0.003283 | 0.9927 | 5 |

<sup>†</sup>One-way ANOVA, Dunnett's multiple comparisons test vs. wild-type.

**Table S7.** Angiotensin II-induced PI(4,5)P2 hydrolysis mediated by HiBit-PLCβ3-CAAX variants.

| construct | $\Delta BRET$ (normalized to wt PLCβ3-CAAX) | | | *n* |
| --- | --- | --- | --- | --- |
|  | mean | S.D. | *P* value<sup>†</sup> |  |
| +GRKct RQ | 1.005 | 0.1263 | - | 7 |
| +GRKct | 0.306 | 0.06239 | - | 7 |
| L40E | 0.5392 | 0.1517 | <0.0001 | 7 |
| R185E | 0.2425 | 0.04493 | <0.0001 | 7 |
| K183E | 0.47 | 0.09543 | <0.0001 | 7 |

<sup>†</sup>One-way ANOVA, Dunnett's multiple comparisons test vs. wild-type.

